# Provenance Legacies Override Species Effects in Shaping Oak Rhizosphere Microbiomes and Metabolomes

**DOI:** 10.1101/2025.11.06.686906

**Authors:** Sebastian Bibinger, Tetyana Nosenko, Prasath Balaji Sivaprakasam Padmanaban, Stefanie Schulz, Hilke Schroeder, Birgit Kersten, Ina Zimmer, Franz Buegger, Michael Schloter, Jörg-Peter Schnitzler

## Abstract

As climate change drives more frequent and intense drought-heat extremes, selecting drought-tolerant trees is crucial for future forest resilience. However, the role of tree-microbial associations for this key trait remains largely unclear. In this study, we investigated how geographic origin, species identity, and intrinsic water-use efficiency (iWUE) shape the rhizosphere microbiome and root-rhizosphere metabolome of pedunculate (*Quercus robur*) and sessile (*Q. petraea*) oaks. In a six-year common garden experiment, we analyzed trees from both species, each grown from seeds from two distinct geographic origins, the upper Rhine basin (URB) and the north-east German lowlands (NGL), differing in water availability, using 16S and ITS rRNA gene based metabarcoding and untargeted metabolomics. We found a consistent legacy effect of seed origin on the composition of the prokaryotic rhizosphere microbiome and the metabolome, whereas tree species had no significant impact. The bacterial family *Pseudonocardiaceae* was enriched at trees from the drier origin NGL, while *Blastocatellaceae* and *Micromonosporaceae* were positively associated with iWUE across samples. Higher iWUE was furthermore significantly correlated with lower prokaryotic diversity and shifts in β-diversity, thereby linking a drought-adaptive host trait to the assembly of the belowground environment. Ellagic acid, a plant derived polyphenol associated with drought tolerance, was enriched in the drier origin NGL and linked to several prokaryotic taxa in correlation networks. The rhizosphere fungal community, however, was largely unaffected by origin or species. Solely fungal community evenness declined with increasing iWUE. Together, our findings suggest that ecotypic adaptation linked to origin can outweigh the effect of species-level traits in shaping the oak rhizosphere microbiome and metabolome. These findings emphasize that provenance-driven ecotypic adaptation can strongly influence plant–microbe interactions and underscore the need for provenance-aware selection and microbiome-informed assisted migration as strategies to strengthen forest drought resilience under global climate change.

## Introduction

Forests cover approximately 30% of Earth’s land area (Hansen et al., 2013) and act as biodiversity hub as well as major global carbon sink. Currently, forests are transforming mainly through the effects of climate change (Allen et al., 2010). The speed of climate change is causing a mismatch between local tree genotypes and their environments, forcing species to shift their geographic ranges (I. C. Chen et al., 2011; Loarie et al., 2009). To counteract this disequilibrium, strategies such as the assisted migration of trees are being considered to align populations with future climates (Aitken & Bemmels, 2016; Zou et al., 2024).

Although climate warming can promote longer growing seasons and CO_2_ fertilization, it also intensifies water scarcity and leads to increased occurrence of combined drought–heat events (Manning et al., 2019; Mukherjee & Mishra, 2021). This poses a significant threat to the health and stability of established forest ecosystems (Thom et al., 2023). Consequently, drought tolerance is a key trait in predicting tree survival in future climates (Allen et al., 2010). This complex trait involves root and leaf morphology, vascular anatomy, and the ability to rapidly acclimatise to a water deficit through processes such as solute accumulation, stomatal regulation, and partial defoliation, as well as other morphological, physiological, and molecular adjustments (Pfenninger et al., 2024).

*Quercus robur* and *Quercus petraea* are dominant oak species of Europe, accounting for more than 9% of forest trees in Germany (Blickensdörfer et al., 2024; Thünen-Institut, 2022). While some oak species are considered to be drought-tolerant and may outperform other European tree species under projected warming conditions (Scharnweber et al., 2011), both provenance and species traits shape their responses to drought (Bruschi, 2010; Arend et al., 2011; Kuster et al., 2013; Rabarijaona et al., 2022). Compared to *Q. robur, Q. petraea* tends to occupy drier, well-drained sites and shows higher intrinsic water use efficiency (iWUE) (Nosenko et al., 2025; Ponton et al., 2001, 2002).

iWUE, as estimated by the δ^13^C value, is an index of carbon gain relative to water loss and can serve as a proxy for drought tolerance (Farquhar et al., 1989; Nosenko et al., 2025). Greater stomatal responsiveness to declining humidity has also been reported in *Q. robur* (Gieger & Thomas, 2005). Across German oak populations, differences in δ^13^C correlate with climate/soil water availability at their origin, particularly under extreme drought, reflecting both constitutive iWUE differences and plasticity (Nosenko et al., 2025). This suggests a genetic basis for adaptation and aligns with evidence of a small number of large-effect genes with contrasting expression patterns between high- and low-iWUE oak genotypes (Brendel et al., 2008; Le Provost et al., 2022).

Belowground tree-microbiota interactions may contribute to the mediation of drought response. Generally, the rhizosphere microbiome, comprising bacteria, archaea, fungi, and protists, plays a pivotal role in plant nutrition, disease resistance, and abiotic stress tolerance (Bulgarelli et al., 2013; Trivedi et al., 2020). It can enhance drought tolerance by improving water and nutrient acquisition, modulating hormonal signalling, altering root system architecture, and priming plant stress responses (Augé, 2001; Carter et al., 2023; Pereira et al., 2020; Rolli et al., 2015; Vurukonda et al., 2016). During drought, microbial communities often shift toward stress-tolerant taxa or into drought-resilient mycorrhizal assemblages (Lekberg et al., 2012; Naylor & Coleman-Derr, 2018).

The rhizosphere microbiome assembly is shaped by plant-controlled exudation patterns (Koprivova & Kopriva, 2022; X. Wang et al., 2024), and varies among species (Ishida et al., 2007; B. Wang & Sugiyama, 2020) and even within genotypes of the same species (Whitham et al., 2012). Genotype effects on associated microbiomes have been observed in annual and perennial species (Brown et al., 2020; da Costa et al., 2022; Schlaeppi et al., 2014; Simonin et al., 2020; Wagner et al., 2016), including trees, for example, *Populus spp*., *Picea* spp., and *Pinus radiata* (Bonito et al., 2019; Gallart et al., 2018; Lamit et al., 2016; Y. Li et al., 2018; Velmala et al., 2013). Furthermore, provenance-related ‘origin legacy’ effect, a lasting imprint of the plant origin on their microbiome, were reported to shape root-associated microbiomes in *Pinus sylvestris, Betula pendula*, and *Fokienia hodginsii* (Färkkilä et al., 2023; H. L. Liu et al., 2024; Maitra et al., 2024). Notably, seed-borne (vertical) transmission can introduce microbiota to the next generation (Abdelfattah et al., 2021; Seitz et al., 2022), potentially interacting with horizontal recruitment from the soil. In fact, seed microbiomes are increasingly recognised as a primary source of inoculum for plant holobionts (Shade et al., 2017; Simonin et al., 2022; U’Ren & Zimmerman, 2021).

Nevertheless, the links between traits and the microbiome that underlie drought adaptation in long-lived trees have not yet been sufficiently explored. Due to their perennial nature, genotypic effects in trees are likely to be more persistent and cumulative than in annual plants, where such influences may be reset each season. A tree’s geographic origin might therefore have a long-lasting impact on the rhizosphere microbial composition and chemistry, on the one hand by genetic adaptation and on the other hand by vertical transmission of microbes of important traits, for example related to drought tolerance. Furthermore, interactions between drought-adaptive traits of the tree, such as intrinsic water-use efficiency (iWUE), and the diversity or composition of rhizosphere microbiomes are largely unknown.

In this study, we examine how the origin (differing in mean annual precipitation) of the tree (ecotype), species identity, and water use efficiency influence the rhizosphere microbiome and metabolome of *Q. robur* and *Q. petraea* trees grown from seeds in a common soil for over six years. Using δ^13^C-based leaf iWUE alongside untargeted root-rhizosphere metabolomics and amplicon sequencing of rhizosphere fungi and prokaryotes, we evaluate two hypotheses:

1. The oak rhizosphere retains origin-specific signatures in fungal and prokaryotic microbiomes and metabolomes even after several years of growth in a shared substrate and environment.
2. Variation in iWUE, reflecting the capacity of a tree to adapt to different environmental conditions, is linked to the associated rhizosphere microbiome composition.

## 1. Materials and Methods

### 2.1 Experimental setup

For this study, two-year-old plants from two tree species, *Quercus robur* (L.) and *Q. petraea* (Matt.) Liebl were bought from a nursery in 2019. The seeds used by the nursery to raise the young trees originated from two forest sites located in the upper Rhine basin (URB; 49.014° N, 8.156° E) and the northeast German lowlands (NGL; 54.144° N, 13.072° E), respectively.

Climate information and climate-derived soil parameters were obtained from the DWD Open Data Portal (https://opendata.dwd.de) as described in (Nosenko et al., 2025). Moisture deficit (MD) was calculated as the difference between mean annual evaporation over grass and loamy sand (VGLS) and mean annual precipitation.

The full origin descriptions can be found in the supplementary material (Supplementary Table S1). The two regions differ in their average annual precipitation (803 mm and 657 mm at URB and NGL, respectively, soil water holding capacity measured within the effective rooting depth of up to 1 m during the vegetation period (307.15 mm and 254.74 mm in URB and NGL, respectively), and moisture deficit (−419.3 and –292.0 in URB and NGL, respectively) (Nosenko et al., 2025).

Seeds from the two origins were grown in a common substrate at a tree nursery (Schrader Pflanzen Handelsgesellschaft GmbH & Co.; Kölln-Reisiek, Germany). In March 2019, 178 bare rooted seedlings of *Q. robur* and *Q. petraea* (79 and 99 from URB and NGL, respectively) were planted to 15-liter pots filled with standard forest soil mixture (Supplementary Table S2) and arranged in a randomized block design in a common garden setup and maintained under mesic conditions with supplemental irrigation at the outdoor area of the Thünen Institute of Forest Genetics, Grosshansdorf, Germany. In the winter of 2021, all seedlings were repotted into 15-liter pots using the same soil substrate. To differentiate between *Q. robur* and *Q. petraea* and exclude hybrid genotypes from the study, molecular marker analysis was conducted on all trees in the common garden as described before (Schroeder & Kersten, 2023). For the root microbiome analysis, 36 oaks (18 trees from species *Q. robur* and *Q. petraea*, respectively) were selected from two origins (20 from NGL and 16 from URB) (Figure 1). The selection covered iWUE values from 57.7 to 116.4. Root samples were collected in February 2023 from the six-year-old plants. Plants were removed from their pots, and roots were sampled from the lower third of the root ball using scissors (Supplementary Figure S1). The cuttings, with soil still attached, were placed in aluminium bags, immediately flash-frozen in liquid nitrogen and stored at −70°C until further processing. In 2022, tree height and basal stem diameter were comparable across all origins and species.

**Figure 1.**
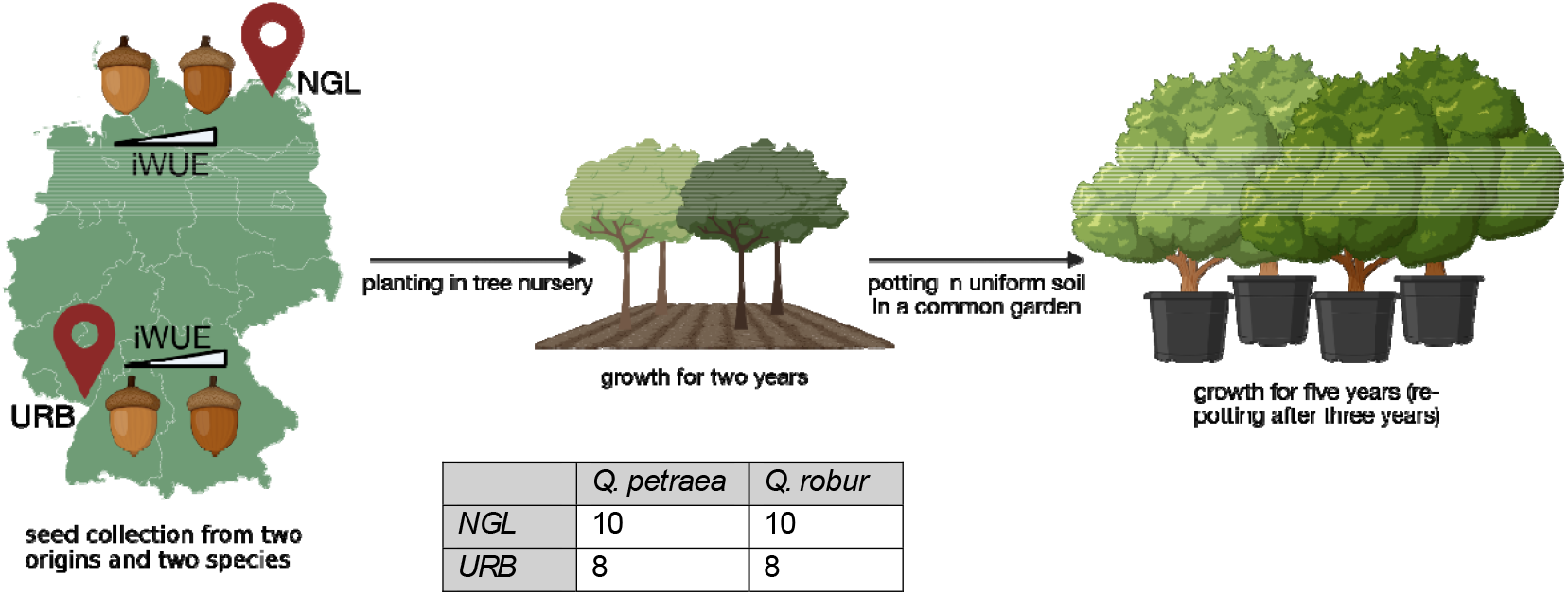
Graphical representation of experimental setup. Seeds were collected at two origins, upper Rhine basin (URB) and north German lowland (NGL). Seeds were then grown for two years in a tree nursery before planting in a common garden. After additional four years of growth, rhizosphere samples were collected in 2023. Intrinsic water use efficiency was determined in 2020. Map lines delineate study areas and do not necessarily depict accepted national boundaries. Figure created with BioRender.com.

### 2.2 iWUE determination

We used intrinsic water-use efficiency (iWUE) as a proxy for the intrinsic tree’s drought tolerance. iWUE was estimated based on the stable carbon isotope ratio (δ^13^C). ^13^C stable isotope analysis was performed in soluble and solid fractions of leaf tissue collected in 2020 from each of the 178 seedlings using an Isotope Ratio Mass Spectrometer (IRMS; delta V Advantage, Thermo Fisher, Dreieich, Germany) and an Elemental Analyzer Euro EA (Eurovector, Milano, Italy) following established procedures (Simon et al., 2013; Nosenko et al., 2025). δ^13^C values are expressed relative to the international Vienna Pee Dee Belemnite (VPDB) standard:

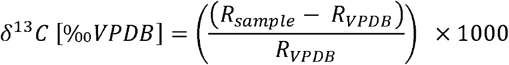

where R_sample_ is the ^13^C/^12^C ratio of the sample and R_VPDB_ = 0.0111802 (Werner & Brand, 2001). iWUE values were then calculated from δ^13^C in the soluble fractions of leaf extracts using the R package isocalcR (Mathias & Hudiburg, 2022). For that, mean temperatures for the period of leaf development (16.04–15.06.2020) were calculated from the daily temperature information of the German Weather Service station 1975 (Opendata.Dwd.de). Atmospheric carbon isotope ratios (δ^13^Catm) were obtained from the atmospheric CO2 and δ^13^Catm database (Belmecheri & Lavergne, 2020).

### 2.3 Untargeted root metabolomics

Metabolite mass feature profiles (from now metabolites) of the root-rhizosphere samples were generated following established procedures (Ghirardo et al., 2020; Hemmler et al., 2018). The analysis was performed using an ultra-performance liquid chromatograph (Ultimate 3000RS, Thermo Fisher, USA) coupled to an ultra-high-resolution quadrupole/time-of-flight mass spectrometer (Impact II QqToF, Bruker Daltonics, Germany). The mass spectrometer was fitted with an Apollo II electrospray ionisation (ESI) source. Two distinct chromatographic separations were applied to each sample: a reverse-phase (RPLC) run in negative ionization mode and a hydrophilic interaction (HILIC) run in positive ionization mode.

Detailed information on the chromatographic gradients, instrument settings, and parameters for mass spectrometry can be found in the publication by Nosenko et al. (Nosenko et al., 2025).

The raw data was processed and analyzed using Metaboscape software (version 4.0, Bruker). For the initial feature finding, parameters were set to a minimum intensity of 1000 counts per spectrum and a peak length spanning at least 7 spectra. This was followed by a recursive feature extraction step, which required a feature to have a minimum length of 6 spectra and be present in at least 2 out of 5 samples to be retained. The workflow also included essential post-acquisition steps such as peak picking, alignment, isotope filtering, and grouping of features via peak-area correlations. To enhance feature identification, ionization settings for both positive and negative modes were specifically configured by defining primary, seed, and common ions.

Metabolite annotation was performed using a hierarchical approach. High-confidence identification was achieved by matching experimental MS/MS spectra against multiple databases, including HMDB, MoNa, and the Vaniya/Fiehn Natural Products Library. For mass features without a library match, a putative annotation was assigned using the software’s “smart formula” generation, which relies on elemental ratios and accurate mass (m/z). While these formulas were subsequently verified against an in-house R-based script and metabolite database (Bertić et al., 2021), only the chemical formula itself was used in subsequent steps to avoid potential misidentification.

Following annotation, the chemical classification of significant features was carried out using the Multidimensional Stoichiometric Compound Classification (MSCC) method. This approach categorizes compounds based on their elemental composition and key ratios, including C, N, H, O, P, S, O:C, N:C, H:C, P:C, and N:P (Rivas-Ubach et al., 2018).

Finally, the data was normalized by subtracting blank signals, and any remaining missing values were imputed with random numbers below the detection limit (range 1-800). This fully processed dataset served as the foundation for all subsequent statistical analyses.

### 2.4 Microbiome analysis

DNA was extracted from approximately 0.5 g of root-free rhizosphere soil using the NucleoSpin Soil kit (Macherey-Nagel, Germany) with buffer SL1 and enhancer SX following the manufacturer’s instructions. For this purpose, bead beating was performed using a Precellys 24 tissue homogenizer (Bertin Instruments, France). Quality of the extracted DNA was assessed photometrically by a NanoDrop 1000 spectrophotometer (ThermoFisher Scientific, USA), and DNA quantity was determined using the Quant-iT PicoGreen dsDNA Assay Kit (Thermo Fisher Scientific, USA) with a Spark® Multimode Microplate Reader (Tecan Trading AG, Switzerland).

As recommended by the Earth Microbiome Project (earthmicrobiome.ucsd.edu), we used the primer pair 515FB-806RB (Apprill et al., 2015; Parada et al., 2016) for amplification of the V4 region of the 16S rRNA gene (from now 16S) and primers ITS3/ITS4 (White et al., 1990) for the amplification of the fungal ribosomal RNA gene region (from now ITS).

The PCR for both target regions was performed in a 25 µl total volume, consisting of 5 ng of template DNA, 12.5 µl NEBNext high-fidelity polymerase (New England Biolabs, USA), 0.3 pmol per primer, and 2.5 µl of 3% BSA. PCR conditions were as follows: 1 min at 98°C; 25 cycles (16S) and 28 cycles (ITS) of 10 s at 98°C, 30 s at 55°C, and 30 s at 72°C; 5 min at 72°C. To confirm successful amplification, 1% (w/v) agarose gel electrophoresis was used. Amplified DNA was then purified using MagSI NGSprep Plus Beads (Magtivio, Netherlands) at a ratio of 0.8:1 beads/PCR product. DNA quantity was again determined by Quant-iT PicoGreen dsDNA Assay Kit with a Spark® Multimode Microplate Reader.

For both 16S and ITS, sequencing indices were added to the PCR products by PCR using the Nextera XT Index Kit v2 (Illumina, Inc., USA) in a total volume of 25 µl, consisting of 10 ng template DNA, 12.5 µl NEBNext high-fidelity polymerase, and 2.5 µl of each indexing primer. The PCR conditions were as follows: 30 s at 98°C; 8 cycles with 10 s at 98°C, 30 s at 55°C, 30 s at 72°C; 5 min at 72°C. PCR products were again purified with MagSI NGSprep Plus Beads. The final DNA quantification was performed through capillary electrophoresis using the 5200 Fragment Analyzer (Agilent, USA). Samples were then pooled equimolarly at 4 nM and sequenced on an Illumina MiSeq using the Reagent Kit v3 (Illumina, Inc., USA). In total, 3,935,957 and 2,639,845 reads were obtained for 16S and ITS sequencing, respectively.

Sequencing adapters were removed from demultiplexed raw sequences using *CutAdapt* v3.5 (Martin, 2011). The raw sequencing data can be found in the NCBI Sequence Read Archive (SRA) under accession number PRJNA1354844 (Bibinger et al., 2025a). Read processing was performed using *DADA2* v.1.34.0 (Callahan et al., 2016) and R v4.4.2 (R Core Team, 2023) in R Studio v2023.09.1 (RStudio Team, 2020). 16S reads were trimmed at 260 bp (forward) and 200 bp (reverse). ITS reads were trimmed at 250 bp (forward) and 209 bp (reverse) and quality filtered using “maxEE = (2,2)”. Resulting ASV sequences were taxonomically classified using the SILVA database v138.1 (Quast et al., 2013) for 16S sequences and UNITE database v10.0 (Abarenkov et al., 2024). Mitochondrial, chloroplast, and singleton reads were excluded from further analysis. Decontamination was carried out using *Decontam* v1.22.0 (Davis et al., 2017). Appropriate sequencing depth was assessed through rarefaction curves using the vegan package v.2.6.-10 (Oksanen et al., 2025). Detailed information about specific read numbers can be found in Supplement Tables S3 (16S) and S4 (ITS). Samples were normalized using cumulative sum scaling (CSS) from the package *metagMisc* v.0.5.0 (Mikryukov, 2025). Phylogenetic trees were computed using *phangorn* v.2.12.1 (Schliep, 2011) and *IQ-TREE* 2 v.2.4.0 (Minh et al., 2020). Fungal guild assignment was performed using the FUNGuild database (Nguyen et al., 2016) and the *FUNGuildR* v.0.3.0 package (Furneaux & Song, 2020).

### 2.5 Statistical analysis

All subsequent statistical analyses were performed with *R v4*.*4*.*2* (R Core Team, 2023) and RStudio v2023.09.1 (RStudio Team, 2020). Unless specified differently, analyses were performed using *phyloseq* v.1.50.0. Data was visualized using *ggplot* v.3.5.1 (Wickham, 2016), viridis v.0.6.5 (Garnier et al., 2024), and *ggpubr* v.0.6.0 (Kassambara, 2023).

ITS and 16S Shannon diversity and ASV richness were calculated using the phyloseq function “estimate_richness()”. To determine ITS and 16S evenness, the “evenness()” function from the package microbiome v.1.28.0 (Lahti & Shetty, 2017) was used. Metabolome diversity was determined as microbiome diversity, but metabolite reads below the detection limit (0-800 reads) were set to zero beforehand. Significance of differences in diversity indices and metabolite abundances were tested using ANOVA or Sheirer-Ray-Hare test. ITS and 16S community composition was visualized in stacked bar plots using the function “trans_abund()$plot_bar” from *microeco* v.1.13.0 (C. Liu et al., 2021).NMDS and PERMANOVA were performed using the “metaMDS()” and “adonis2()” functions from the package vegan v.2.6-10 (Oksanen et al., 2025). To ensure that differences in group dispersion did not confound PERMANOVA results, we used “betadisper()” to confirm homogeneity of variances beforehand. Differentially abundant families between origins, species, and along iWUE were identified using the DESeq2 v.1.48.1 package (Love et al., 2014) with false discovery rate correction. Family was chosen as the taxonomic level, since on the ASV level, no significant results were found. β-NTI values for community assembly processes were determined using the comdistnt() function from the picante v.1.8.2 package (Kembel et al., 2010).

OPLS-DA was performed on metabolite data using the R package ropls v.1.40.0 (Thevenot et al., 2015). Metabolite data was log10-transformed and Pareto-scaled beforehand using the package IMIFA v.2.2.0 (Murphy et al., 2020). Correlation of ASVs with metabolites was performed through the “rcorr()” function from Hmisc v.5.2-3 (Harrell Jr, 2025). Initially, metabolites were selected based on an OPLS-DA VIP score greater than 1.0, a p-value less than 0.05, and an absolute log2 fold change greater than 1.0. The selected metabolite data was then log10-transformed and Pareto-scaled.

Microbiome data was transformed using a center log ratio (clr) transformation from the package compositions v.2.0.8 (van den Boogaart et al., 2024). Correlations with Spearman’s rho > 0.6 were used to generate a correlation network using igraph v.2.1.4 (Csardi & Nepusz, 2006).

The R scripts for analysis were formatted, refactored, and debugged with assistance from the large-language model Gemini (Google LLC). AI-generated code was always manually reviewed, validated, and adapted. Processed data for final statistical analysis can be found in the Zenodo data archive (Bibinger et al., 2025b), and the R code used for all statistical analyses and figure generation is available in a separate archive (Bibinger et al, 2025c).

## 3. Results

### 3.1 Origin-based legacies persist after six years in common soil

#### 3.1.1 Prokaryotic α-diversity and β-diversity show origin effects

We retrieved an average of 65,555 high-quality reads per sample from 16S rRNA gene metabarcoding. This was sufficient for covering the present biodiversity, as shown by rarefaction curves (Supplementary Table S3 and Figure S2A). Across all 36 samples, we obtained 10,203 unique ASVs, of which 10,177 were assigned to bacteria and 26 were assigned to archaea. The bacterial community was dominated by members of the phyla *Pseudomonadota*, with an average relative abundance of 35.5% (±2.2% SD), *Actinomycetota* (16.5% ± 3.1%), *Acidobacteriota* (12.9% ± 1.4%), and *Chloroflexota* (9.4% ± 2.1%) throughout the samples. The most prominent families included *Xanthobacteraceae* (5.2% ± 0.7%), *Nitrosomonadaceae* (3.3% ± 0.9%), and *Pirellulaceae* (3.0% ± 0.6%) (Supplementary Figures S3A and S3B).

ASV richness, Shannon index, and Pielou’s evenness metric were determined to investigate the microbial alpha diversity (Figure 2A). There were no differences in ASV richness or Shannon diversity between the two species. However, differences in diversity were observed between tree origins. Prokaryotic microbiomes from origin URB showed higher ASV richness (mean ± SD: 1603 ± 343) and Shannon diversity (6.73 ± 0.19) than microbiomes from NGL (Observed: 1311 ± 216; Shannon: 6.54 ± 0.15) at statistical significance (p_observed_ = 0.001; p_shannon_ = 0.004). If splitting the data by tree species, a subsequent Wilcoxon test confirmed the statistical significance of origin, only for prokaryotic Shannon diversity within *Q. robur*, which showed a significant difference between NGL and URB (p = 0.009). Prokaryotic Pielou’s evenness was generally comparable between origins and species.

**Figure 2.**
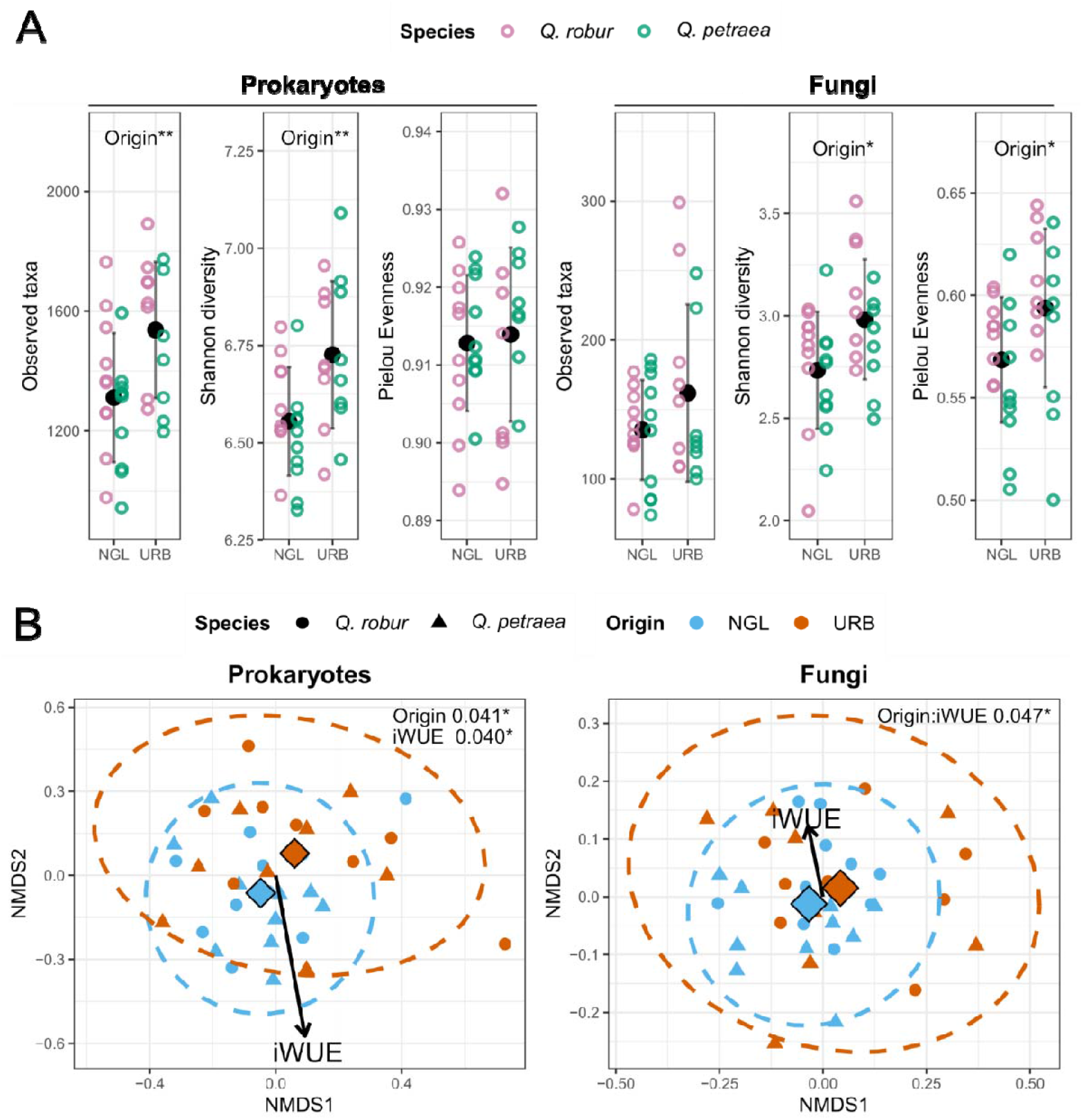
Effects of origin and intrinsic water use efficiency (iWUE) on prokaryotic and fungal microbiome diversity and structure. **(A)** Alpha diversity found in the rhizosphere split by origin and species. Richness, Shannon diversity, and Pielou evenness were used to assess alpha diversity. Significant factors as determined by ANOVA are noted in the plot. For visual clarity, the y-axis was constrained, hiding one outlier in prokaryotic richness and one in fungal evenness. **(B)** NMDS based on Bray-Curtis distances was used to assess β-diversity. The arrows indicate the direction of increasing iWUE. Diamond shapes represent the origin centroids. Significant factors, as determined with PERMANOVA, are shown in the plot. The full results from the PERMANOVA model can be found in the supplement (Supplementary Table S5).

Permutational multivariate analysis of variance (PERMANOVA) based on Bray–Curtis distances revealed compositional differences in the prokaryotic communities of trees originating from URB and NGL (Figure 2B). Although the effect size was small (R^2^ = 0.04), the difference was significant (p = 0.024). No compositional differences in the prokaryotic communities between tree species were detected (p = 0.662). To pinpoint specific taxa driving the observed differences, we performed a differential abundance analysis using DESeq2. While we could not find differentially abundant ASVs, a family-level analysis revealed an accumulative effect which resulted in significant enrichment of *Pseudonocardiaceae* in origin NGL (log2FC = 1.27, p = 0.014, predominately genera *Amycolatopsis* and *Pseudonocardia*), and *Sphingobacteriaceae* (log2FC = 1.68, p = 0.014, predominately genera *Pedobacter* and *Mucilaginibacter*) in origin URB (Supplementary Figure S5A). Consistent with our PERMANOVA results, no differentially abundant families or ASVs were found between the tree species.

#### 3.1.2 Fungal alpha but not beta diversity shows origin effects

From ITS sequencing, we received 35,098 high-quality reads per sample, representing 1,902 unique ASVs, which also covered fungal biodiversity sufficiently (Supplementary Table S4 and Figure S2B).

Fungal community displayed a higher variability in their composition. Dominating phyla were *Ascomycota* (62.8% ± 13.3%) and *Basidiomycota* (31.4% ± 13.1%), together accounting for approximately 94.0% of the total fungal sequences. Highly abundant fungal families, besides unclassified families from the phylum *Ascomycota*, include the ectomycorrhizal family *Sclerodermataceae* (27.1% ± 13.1%), *Gymnoascaceae* (7.8% ± 5.1%), and *Pseudeurotiaceae* (5.8% ± 4.4%) (Supplementary Figures S3C and S3D). We included an assessment of fungal guild assignment using the Funguild database. The most prominent guilds identified were ectomycorrhizae (EMF) and undefined saprotrophs (Supplementary Figure S4). The EMF community was dominated by *Scleroderma areolatum* (76.2%) and *Scleroderma verrucosu* (9.6%).

Fungal ASV richness showed no significant difference between origins or species (Figure 2A). However, NGL fungal communities exhibited significantly lower Shannon diversity than URB communities (2.73 ± 0.29 vs. 2.98 ± 0.29; p = 0.036), which was driven by the same pattern in Pielou’s evenness (0.56 ± 0.04 vs. 0.59 ± 0.04; p = 0.014). Again, subsequent Wilcoxon tests for origin effects within each tree species separately did not confirm any significant differences between origins (Q. robur: p_shannon_ = 0.122, p_pielou_ = 0.266; *Q. petraea:* p_shannon_ = 0.274, p_pielou_ = 0.237).

Fungal composition was not significantly different between tree species or origins (p = 0.493 and 0.077), as determined through PERMANOVA (Figure 2B). However, the interaction of origin and iWUE had a significant effect on fungal microbiome composition (R^2^ = 0.047, p = 0.014). We found no significant impacts of any factors on the ECM subcommunity. Consistent with our PERMANOVA results, no differentially abundant families or ASVs were found between the tree species or origins using DESeq2.

#### 3.1.3 Metabolomes are separated by origin and differ in their class composition

Across all samples, 1,071 metabolites were identified and assigned to six classes: carbohydrates, lipids, amino sugars, nucleotides, proteins, and oxyaromatic compounds. Metabolite richness, Shannon index, and Pielou’s evenness metric were determined to investigate the metabolite alpha diversity (Figure 3A). Metabolite richness was significantly higher in samples from origin NGL than from URB (701 ± 95 and 624 ± 100, ANOVA p = 0.02). Similar results were found for Shannon diversity (5.65 ± 0.10 and 5.53 ± 0.18, p = 0.02). Metabolite evenness, on the other hand, was significantly higher in Q. petraea compared to Q. robur (0.87 ± 0.01 and 0.86 ± 0.01, p = 7e-06).

**Figure 3.**
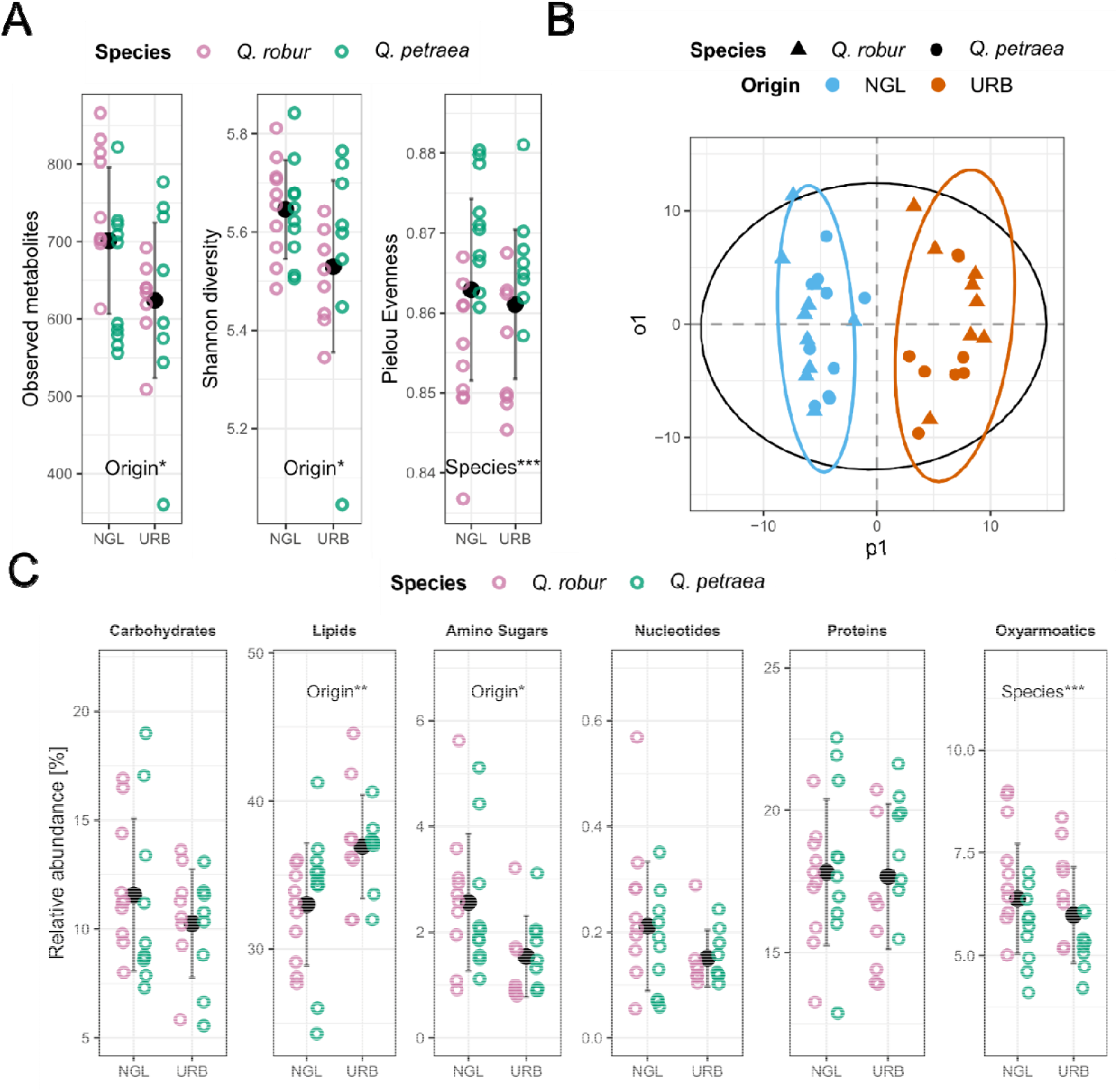
Effects of origin, species, and intrinsic water use efficiency (iWUE) on rhizosphere metabolome composition. **(A)** Metabolome Shannon diversity split by tree origin and species. Significant differences as determined by ANOVA are displayed. **(B)** OPLS-DA scores plot discriminating between metabolomes from URB and NGL. **(C)** Metabolite class relative abundance per species and origin. Significant differences, as found by ANOVA, are marked in the plot.

To investigate compositional differences between the tree origins and species, we performed an Orthogonal Projections to Latent Structures Discriminant Analysis (OPLS-DA). While a model for the factor origin could be constructed, no significant predictive or orthogonal components could be identified for the factor tree species. For the factor origin, OPLS-DA showed a clear separation in metabolites between NGL and URB (Figure 3B). The model accounted for 90.7% of the variation in the response variable (R^2^Y) and 19.3% of the variation in the predictor variables (R^2^X) and showed strong predictive performance (Q2 = 0.648, p = 0.05). A total of 58 metabolites were identified as key drivers of the separation between origins, meeting the criteria of a VIP score > 1, log-fold change (log2FC) > 1, and p-value < 0.05. The abundance of these 58 metabolites and their classification can be found in Supplementary Figure S6 and Table S6. Five key metabolites (two lipids, two proteins, and one unmatched compound) were enriched in origin URB, while 53 metabolites were enriched in origin NGL. The top ten metabolites with the highest VIP score included five lipids, three proteins, and one plant-derived oxyaromatic mass feature classified as ellagic acid. Ellagic acid with a VIP score of 3.49 was enriched in the rhizosphere of NGL with a log2 fold change of 2.69 (p < 0.01).

Concerning metabolite classes, we found comparable carbohydrate abundance between tree species and origins (Figure 3C). For lipids, samples from origin URB showed a significantly higher abundance (36.9% ± 3.5%) than those from NGL (33.0% ± 4.1%), as determined by ANOVA (p = 0.006). An opposite trend was observed for amino sugars, which showed significantly higher abundance in samples from NGL (2.6% ± 1.3%) compared to URB (1.5% ± 0.8%, ANOVA p=0.010). While nucleotide and protein abundance were comparable between species and origins, samples from *Q. robur* contained significantly higher amounts of oxyaromatic compounds (6.9% ± 1.3%) than *Q. petraea* (5.4% ± 0.8%, ANOVA p < 0.001). Overall, 27.9% of the metabolites could not be assigned to a class.

### 3.2 Microbiome–metabolome coupling is limited globally but reveals shared hubs

A Procrustes analysis of the PCA scores was used to assess global congruence between microbiome and metabolome profiles. On the one hand, no significant association was found for either prokaryotes or fungi (prokaryotes: Procrustes r = 0.257, p = 0.16; fungi: r = 0.2544, p = 0.18). On the other hand, there was a significant negative correlation between metabolite and prokaryotic Shannon diversity (rho = −0.46, p = 0.006, Supplementary Figure S7), which did not extend to fungal diversity (rho = −0.08, p = 0.649). To explore specific microbe-metabolite relationships, we conducted a Spearman’s partial correlation analysis between the 58 key metabolites and the members of the prokaryotic and fungal microbiomes (Figure 4).

**Figure 4.**
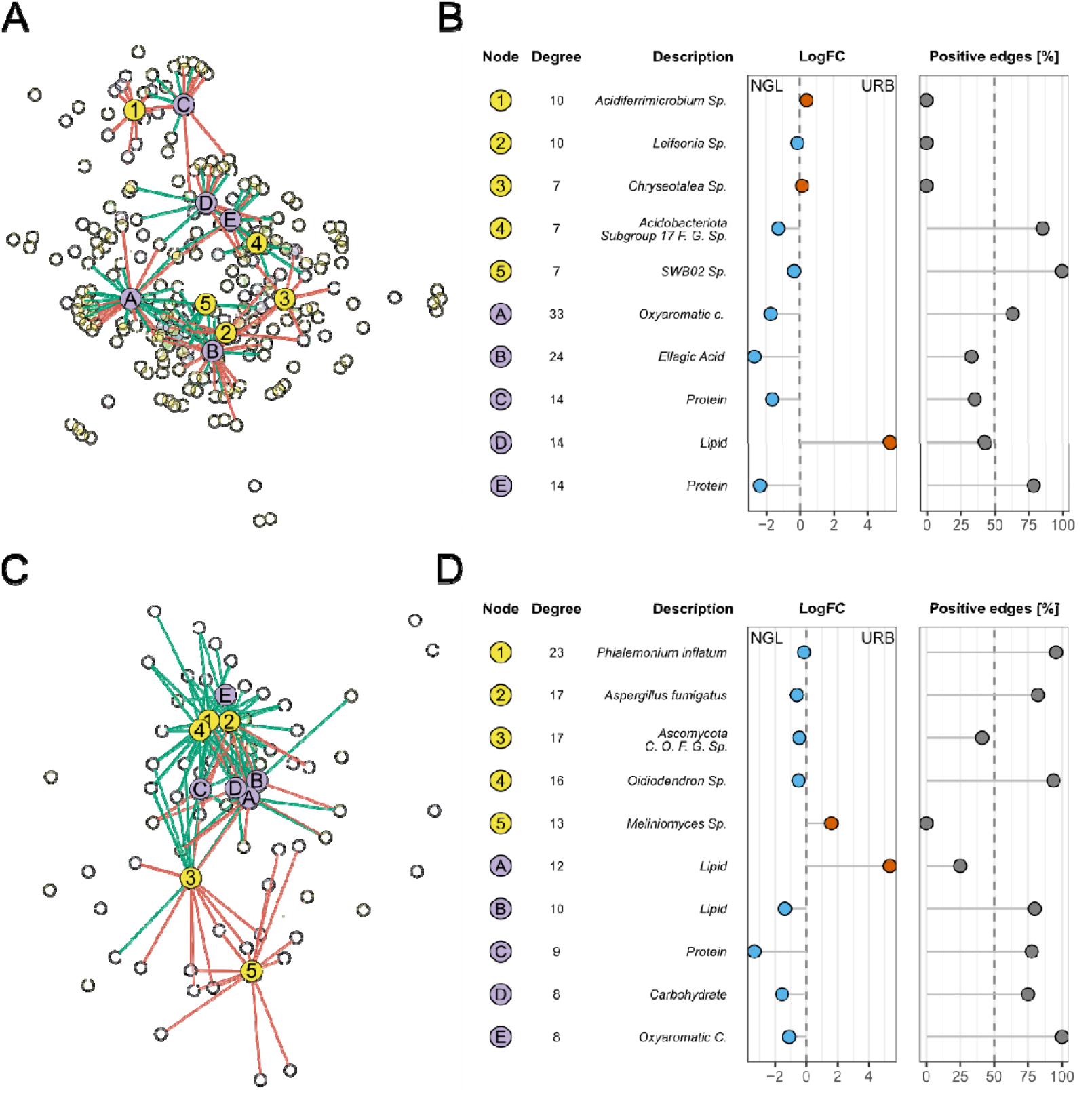
Correlation analysis between key metabolites (purple) and prokaryotic and fungal taxa (yellow). Spearman partial correlation network of prokaryotic (A) and fungal (C) microbiomes and key metabolites. Network edges represent pairwise associations (Spearman’s ρ > 0.4) between microbes and metabolites, but without statistical significance. Characterization of the top 5 connected metabolites and prokaryotic (B) and fungal (D) ASVs. Shown are their degree (number of correlations with correlation coefficient > 0.4), taxonomic classification or assigned metabolite class, their abundance log2 fold change between the origins (coloured by origin of enrichment), and their positive edge (positive correlation) percentage.

Correlation networks were constructed, with edges representing significant correlations (rho > 0.4) between metabolites and prokaryotic (Figure 4A) or fungal ASVs (Figure 4C). The prokaryotic network contained 234 nodes and 408 edges with 55.9% positive edges. The fungal network contained 76 nodes and 208 edges with 67.8% positive edges. Based on those networks, we characterized the nodes (taxa and metabolites) with the highest degree (i.e., number of connections; Figure 4B and 4D). Notably, the top three bacterial taxa with the highest number of strong correlations, *Acidiferrimicrobium* sp. (degree = 10), *Leifsonia* sp. (10), and *Chryseotalea* sp. (degree = 7) showed only negative correlations with metabolites. The other two bacterial taxa, one from the phylum *Acidobacteriota* (degree = 7) and one from the genus SWB0 from the family *Hyphomonadaceae* (degree = 7), showed mostly positive correlations (85.7% and 100%). None of the top connected taxa showed significant differential abundance between the origins.

Notably, ellagic acid was the metabolite with the second-highest degree in the prokaryotic network (24) and was also found within the top ten metabolites with the highest VIP score. It was enriched in origin NGL with a log2 fold change of −2.69 and showed 33.3% positive correlations. The other, not further classified, metabolites with a high degree were one oxyaromatic compound (degree = 33) and two proteins (both degree = 14), which were significantly enriched in origin NGL (log2FC = −1.62 and −2.39) with positive edge percentages of 35.7% and 78.6%, respectively. Conversely, one lipid (degree = 14) that was strongly enriched in origin URB (log2FC = 5.35) with a positive edge percentage of 42.9% was also within the top five connected nodes. This compound was annotated as an iridoid similar to plant-derived valeriotetrate.

For metabolite interactions with fungal ASVs, three of the ASVs with the highest degree, *P. inflatum* (degree = 23), *Aspergillus fumigatus* (degree = 17), and *Oidodendron* Sp. (degree = 16), showed mostly positive correlations with the metabolites (95.7%, 82.3%, and 93.8%). An ASV of phylum *Ascomycota* (degree = 17) and one of genus *Meliniomyces* (degree = 13) showed lower positive edge percentage (41.1% and 0.0%). None of the ASVs showed significant differential abundance between the origins. Four metabolites, one from each of the following classes: lipids, proteins, carbohydrates, and oxyaromatic compounds, were enriched in NGL (log2FC = −1.37, −3.33, −1.55, −1.10) and showed predominantly positive correlations (80.0%, 77.8%, 75.0%, 100%). Once again, there was one URB-enriched lipid in the top five connected metabolites (log2FC = 5.35, degree = 12). This lipid was the same one already found in the prokaryotic network (Node D). It had a positive edge percentage of 25.0% in the fungal network.

### 3.3 iWUE impacts prokaryotic but not fungal composition and diversity

According to the experimental design, where intrinsic water use efficiency (iWUE) was implemented as an independent tree variable across sites, tree origin or species had no significant effect on this parameter (Table 1). Nevertheless, iWUE was slightly higher on average in trees originating from NGL compared to URB (p = 0.078). Since trees were grown in pots and therefore height and diameter could have been technically influenced, we do not interpret these parameters and exclude them from further analysis.

**Table 1.** Intrinsic water use efficiency (iWUE) average values and SD by oak species and origin.

| Species - Origin | iWUE |
| --- | --- |
| *Q. robur* – NGL | 88.3 ± 13.2 |
| *Q. petraea* – NGL | 92.9 ± 16.5 |
| *Q. robur* – URB | 80.5 ± 22.0 |
| *Q. petraea* – URB | 79.91 ± 16.24 |

Spearman’s rank correlation analysis revealed that trees with higher iWUE values had lower prokaryotic diversity in the rhizosphere microbiome. A significantly negative correlation was found between iWUE and both prokaryotic richness (rho = −0.50, p = 0.002) and Shannon diversity (rho = −0.60, p = 0.002, Supplementary Figure S8) across all samples (Figure 5). This correlation was also observed for the NGL and *Q. robur* data subsets (Table 2). PERMANOVA revealed furthermore that iWUE of the tree significantly impacted the prokaryotic community composition (R^2^ = 0.04, p = 0.036) (Figure 2B). Within samples from the origin NGL, we found *Blastocatellaceae* (log2FC = 0.30, p = 0.024) and *Micromonosporaceae* (log2FC = 0.47, p = 0.008) to be positively associated with iWUE (DESeq2 results; Supplementary Figure S5).

**Table 2.** Spearman correlation between microbial richness and iWUE for sample subsets. Shown are un-adjusted p-values.

|  | Prokaryotes |  | Fungi |  |
| --- | --- | --- | --- | --- |
|  | rho | p-value | rho | p-value |
| full | -0.50 | **0.002** | 0.07 | 0.577 |
| NGL | -0.59 | **0.007** | 0.41 | 0.073 |
| URB | -0.18 | 0.498 | -0.11 | 0.688 |
| *Q. robur* | -0.56 | **0.016** | -0.10 | 0.705 |
| *Q. petraea* | -0.43 | 0.076 | 0.26 | 0.291 |

**Figure 5.**
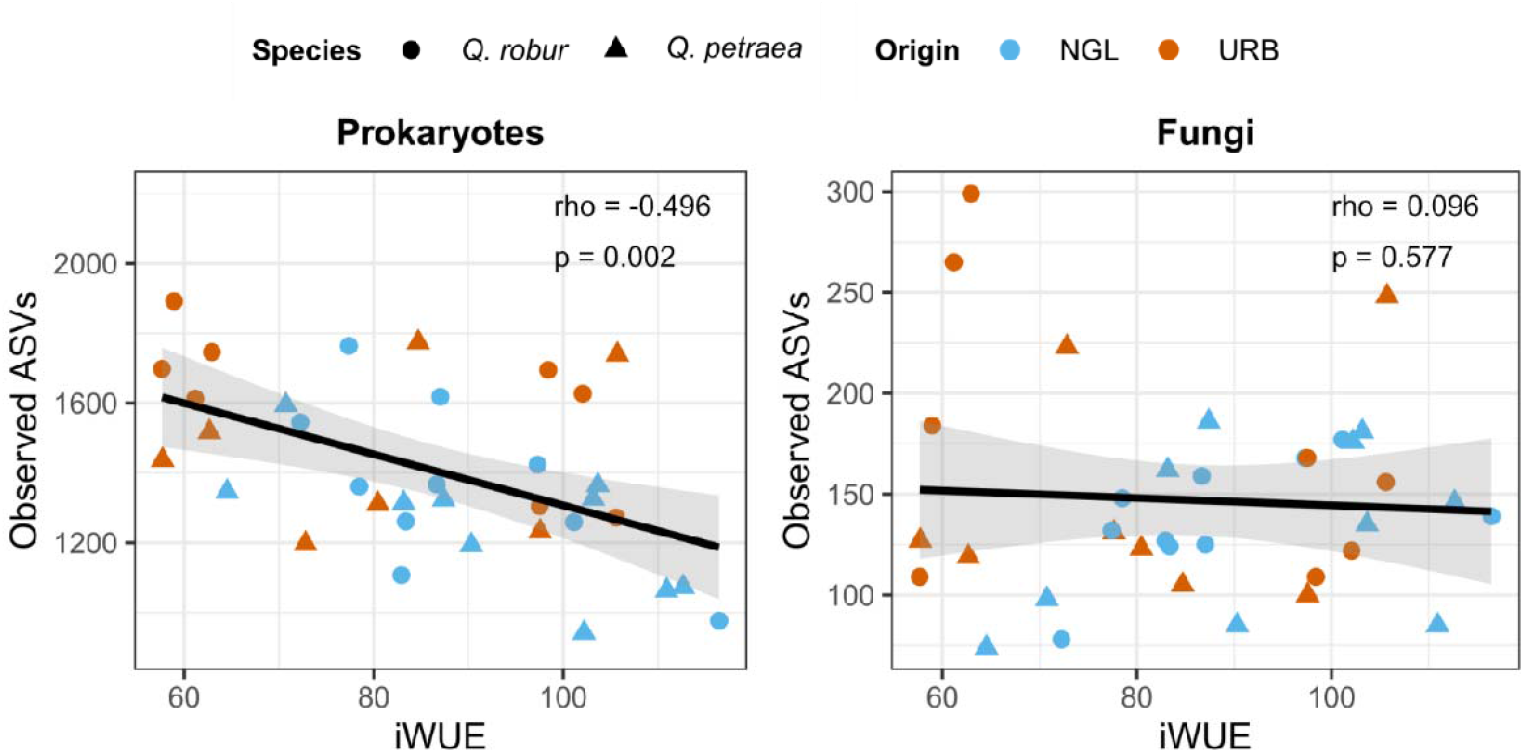
Spearman correlation of microbial richness and intrinsic water use efficiency (iWUE) across species but split between origin. Spearman correlation coefficient and p-values are indicated in the plot. Correlation between fungal evenness and iWUE was not significant.

We did not observe any significant relationship between richness and iWUE for fungal communities (Figure 5). However, fungal evenness showed a significant negative relationship with iWUE (rho = −0.46, p = 0.005). While the fungal community composition did not differ significantly between tree species or seed origins, the interaction between origin and iWUE significantly affected it (R^2^ = 0.05; p = 0.014). Finally, we could not associate fungal taxa or families with iWUE using DESeq2.

## 4. Discussion

In the present study, we found that the origin of the *Q. petraea* and *Q. robur* seeds has a lasting effect in the rhizosphere after years of growth under identical conditions, shaping its prokaryotic microbiome and metabolome. However, we observed neither significant species-specific effects on the oak rhizosphere nor a similar response from the fungal community. Furthermore, independently of origin, higher tree iWUE was significantly associated with lower prokaryotic alpha diversity in the rhizosphere.

### 4.1 Tree origin legacy effect on rhizosphere microbiome and metabolome

The origin of the seeds had a small but significant effect on the composition of the prokaryotic community. The compositional shift was characterized by the increase of the *Pseudonocardiaceae* diversity in the rhizosphere of trees originating from a relatively dry region, NGL, and an increase of *Sphingobacteriaceae* at URB, characterized by a higher precipitation level. The enrichment of *Pseudonocardiaceae* at the drier site is notable; this family, including genera like *Pseudonocardia* and *Amycolatopsis*, is known for its drought tolerance and various plant growth-promoting traits (Borah & Thakur, 2020; Chaiya et al., 2021; Xie et al., 2024). Furthermore, the genus *Pseudonocardia* also includes endophytic species (Qin et al., 2011), which suggests a potential role in the vertical microbiome transmission mentioned later. Furthermore, trees from the origin URB exhibited higher rhizosphere prokaryotic and fungal Shannon diversity. While the higher Shannon index of the prokaryotic community in URB was driven by higher richness, the fungal diversity difference was caused primarily by increased evenness.

The detection of differences in microbiome composition between origins aligns with previous research demonstrating that seed or plant origin can shape tree microbiomes, like prokaryotic endophytes and fungal rhizosphere communities (Färkkilä et al., 2023; Maitra et al., 2024). Our results are consistent with the results of recent work by Liu et al. (H. L. Liu et al., 2024), who identified significant differences in the rhizosphere microbiomes among 11 provenances of *Fokienia hodginsii*. By demonstrating this phenomenon through a long-term experiment with a robust, biologically replicated design, our study corroborates their findings regarding an origin legacy effect.

We observed no differences in the composition of the fungal or prokaryotic communities between *Q. petraea* and *Q. robur* at the species level. This finding was surprising, since rhizosphere microbiome differences between different tree species can be expected to be stronger than between intraspecific ecotypes and are frequently described (Ishida et al., 2007; B. Wang & Sugiyama, 2020). On the other hand, differences in the rhizosphere can be small and hard to detect and are often observed in adult trees (Bonito et al., 2019; Cregger et al., 2018). The eco-geographic adaptation to the region of origin might have overwritten the species-specific effects, especially considering the close relatedness of both species.

To investigate the potential drivers behind the origin of the legacy, we examined the rhizosphere metabolome as a link between the tree phenotype and microbiome composition. The metabolome composition mirrored the pattern that we observed in microbiomes. OPLS-DA separated metabolomes by origins but not by species. Interestingly, the metabolite alpha diversity was higher in trees from NGL, displaying a reverse trend of what was observed in the prokaryotic microbiome. Furthermore, the metabolite evenness was higher in *Q. petraea* than in *Q. robur*. While large environmental surveys often find positive metabolome-microbiome covariation (Shaffer et al., 2022), we detected no global congruence (Procrustes) and found a negative association between prokaryotic and metabolite Shannon diversity. This divergence could reflect targeted exudation under drier conditions, metabolic channeling by subsets of taxa, or ontogenetic exudation traits in juveniles, as further discussed below.

Through exploratory correlation network analysis, we identified plant derived ellagic acid as a potential mediator of drought response. It was significantly enriched in the drier origin NGL and was highly correlated with a range of prokaryotic taxa. Ellagic acid is abundant in oaks (Othón-Díaz et al., 2023). It can alleviate water deficit stress via antioxidant pathways (Debnath et al., 2020; García-Niño & Zazueta, 2015). For example, seed pretreatment with ellagic acid has been shown to enhance drought tolerance in seedlings of *Cicer arietinum* (chickpea) and *Brassica napus* (canola) (El-Soud et al., 2013; Khan et al., 2017). Furthermore, treating maize with ellagic acid effectively reduced drought stress symptoms (Agar et al., 2024). Based on our results, ellagic acid may have another function via interactions with the rhizosphere microbiome. The other notable metabolite, which was strongly enriched in origin URB, was a valeriotetrate-like iridoid, a class of plant derived compounds that are involved in defense against herbivores and pathogens (Biere et al., 2004; Bowers & Stamp, 1993). Among microbes, *Leifsonia*, which is frequently found in the rhizosphere and is implicated in osmotic stress tolerance (Kang et al., 2017; Nordstedt et al., 2021), and *Phialemonium inflatum*, which is reported to promote growth and suppress pathogens (Rivera-Vega et al., 2022; Zhou et al., 2018), were highly connected. As microbial communities shape rhizosphere chemistry themselves, these patterns likely reflect bidirectional feedback between exudation and microbial community structure, rather than unidirectional plant control.

### 4.2 Mechanistic routes for legacy persistence

Origin legacy effects in the rhizosphere microbiome can be mediated in two ways: vertical and horizontal transmission (Gundel et al., 2011; Shade et al., 2017). Vertical transmission, carryover of the microbiome from the mother tree via the seed, was previously seen in several plants, including oaks (Abdelfattah et al., 2021; Fort et al., 2021; Hardoim et al., 2012; Johnston-Monje & Raizada, 2011; Walitang et al., 2019). It is unclear whether the transmitted species could remain on the next generation of plants for up to six years (Vandenkoornhuyse et al., 2015). Horizontal transfer depends on the ability of ecotypes to adapt to their specific habitats. Such adaptation leads to genetic or epigenetic changes that are inherited across generations. Resulting variations in root morphology and exudation between tree ecotypes have been widely reported and could alter the recruitment of a specific microbiome from the bulk soil (H. L. Liu et al., 2024; Luo et al., 2017; Seitz et al., 2022).

Here, it must be noted that the samples were collected from six-year-old trees, which were still in the juvenile stage of development. The root exudation patterns change significantly with tree development phase. Often the total amount of exudates reduces with age, and the focus shifts from nutrient acquisition towards a greater investment in defensive capabilities (Chaparro et al., 2013; M. Chen et al., 2023; Z. Li et al., 2021). The selective pressure exerted by juveniles on their rhizosphere microbiome may be weaker, allowing seed-transmitted communities to persist alongside the progressive horizontal acquisition of microbiomes. Both a strong origin adaptation and a weak exudative specificity at this juvenile stage could explain the lack of differences in microbiomes and metabolomes between species while accounting for origin-specific differences. Moreover, this might also put juvenile trees in a vulnerable position under climate change, being dependent on their parent trait transfer for successful survival during early growth stages (Au et al., 2022).

Ultimately, knowledge of the original seed microbiome and analysis of root exudation (and other phenotypic or genotypic parameters) could help to clarify how the origin legacy effect is mediated in oaks.

### 4.3 Rhizosphere fungal microbiome is less sensitive to plant traits and origin than the prokaryotic microbiome

Although the seed origin and iWUE were tracked in prokaryotic microbiomes, the fungal rhizosphere showed no comparable effects of origin, species or iWUE, and displayed fewer metabolite interactions. This result also held for ectomycorrhizal fungi (primarily the typical oak symbionts from genus *Scleroderma* (Bzdyk et al., 2018), despite the frequent host specificity reported for this guild (Ishida et al., 2007). This is consistent with reports that the composition of ECM communities is often more influenced by environmental properties than by host traits (Downie et al., 2021; Karliński et al., 2013). As the soil conditions in our experiment were identical across trees, large differences in fungal composition were not anticipated.

Furthermore, several studies demonstrated stronger effects of host traits on bacteria than on fungi (Gaete et al., 2021; Merino-Martín et al., 2020; Song et al., 2025; Ye et al., 2021). Finally, fungi are generally more dispersal-limited than prokaryotes (Song et al., 2025), which in turn contributes to higher inter-sample heterogeneity as rhizosphere communities assemble from a more localized soil environment before (Collins et al., 2018; Larsen et al., 2065; Merino-Martín et al., 2020; Zhang et al., 2021). This increased variability can reduce the specificity of the fungal microbiome to its host.

### 4.4 Diversity of the rhizosphere microbial community is associated with tree intrinsic water-use efficiency

In our analysis, intrinsic water-use efficiency (iWUE) was a significant predictor of prokaryotic community composition (PERMANOVA, R^2^ = 4.0%) and was negatively correlated with prokaryotic alpha diversity. Thus, trees with higher iWUE harbored less diverse prokaryotic communities, regardless of the seed origin or species.

The rhizosphere microbiome can improve the host drought tolerance via alteration of root morphology, exudation profiles, or stomatal conductance (Augé et al., 2015; Carter et al., 2023; Kannenberg & Phillips, 2017). The specific rhizosphere microbiome can therefore be seen as a dimension of drought tolerance adaptation (Ben Zineb et al., 2024; Ulrich et al., 2019). Consequently, Gaete *et al*. (Gaete et al., 2021) reported that the prokaryotic microbiomes of drought-tolerant and drought-susceptible tomato plants differ in their alpha diversity, beta diversity, and interaction network complexity. Contrary to our results, these differences were dependent on drought rather than being apparent under normal irrigation conditions.

Direct evidence linking iWUE and rhizosphere diversity remains limited and is likely to be context-dependent (Naylor & Coleman-Derr, 2018). We hypothesize that adaptation to drier conditions increases the specificity of exudate-mediated selection, yielding more specialized and less diverse microbiomes. The stronger correlation between iWUE and richness at NGL supports this hypothesis. Within the trees originating from NGL, two prokaryotic families, *Blastocatellaceae* and *Micromonosporaceae*, were positively associated with iWUE. Members of both families have been previously found in plant rhizosphere. Although, they had not yet been linked to drought tolerance or growth promotion in plants (Abdullaeva et al., 2024; Ling et al., 2022; Long et al., 2024; Sun et al., 2019), their affiliation with the stress-resilient groups *Acidobacteriota* and *Actinobacteria* (Lavallee et al., 2024) makes them potential contributors to plant drought responses. We predict that the compositional differences between the rhizospheres of oak trees of different origins may become more pronounced under drought stress.

## 5. Conclusion and implications

Our data demonstrates the importance of ecotypic adaptation for the structure of the rhizosphere environment of oak trees. The origin of seeds leaves a long-lasting imprint on the metabolome and prokaryotic diversity and composition of oak rhizospheres, whereas species effects are minimal. Leaf intrinsic water-use efficiency (iWUE), a trait associated with drought adaptation, covaries with these below-ground features independently of the tree origin. This suggests the existence of a trait-microbiome axis that is relevant to drought resilience. The lack of a strong response from the fungal community indicates that prokaryotic and fungal communities in the rhizosphere are shaped by different ecological drivers.

In the face of climate change-driving rapid range shifts, assisted migration is a key forest adaptation strategy.

Given that assisted migration alone cannot completely prevent declines in forest ecosystem services, selecting trees based on origin or functional traits like iWUE may be a more effective strategy than relying on species-level traits alone. As the effects observed in our experiment are modest yet consistent across biomes, they reveal parameters that can be used in process-based models to couple soil microbes with plant water/carbon exchange. Pairing provenance selection with a microbiome-aware approach offers a practical solution to the challenges of assisted migration and forest restoration in the context of global change.

While our conclusions are constrained by the juvenile stage of the oaks, the common-garden pot context, and the correlative nature of the study, they motivate targeted future research. To elucidate the mechanisms underlying the legacy effects and verify their functional relevance for drought tolerance, causal tests should be conducted, such as reciprocal microbiome transfers, mechanistic analyses of root exudation dynamics, and long-term ontogenetic tracking.

Together, these efforts will refine our understanding of how provenance and functional traits shape oak– microbiome systems, and inform provenance-aware, microbiome-informed strategies to enhance forest resilience.

## Acknowledgements

This work has been supported by the grants 2220WK09A4 and 2220WK09B4 from the German Federal Ministry of Food and Agriculture (BMEL), Federal Ministry for the Environment, Nature Conservation and Nuclear Safety (BMU) in the frame of the Waldklimafonds Program. This study was additionally supported by the German Research Foundation (DFG) (Grant 457330647), as part of the Research Unit 5315.

## Competing interests

The authors declare no conflict of interest.

## Author contributions

TN, HS, MS, and JPS designed this study; TN, HS, and BK conducted experiments; SB, TN, IZ, SS, and PBSB analysed and interpreted the data; BK, HS; and JPS administered the project; SB wrote the manuscript, with input from all authors. All authors read and approved the final version of the manuscript.

## Data availability

The raw data (16S and ITS sequences) are available in the NCBI Sequence Read Archive (SRA) under accession number PRJNA1354844 (Bibinger *et al*., 2025a) All processed datasets, including phyloseq objects, metabolite abundance matrices, and metadata required to reproduce the analysis, are archived on Zenodo (Bibinger *et al*., 2025b; https://doi.org/10.5281/zenodo.17514228). The R code used for sequence processing, statistical analysis, and figure generation is publicly available on GitHub at https://github.com/SebastianBibinger/Oak_provenance and permanently archived on Zenodo (Bibinger et al., 2025c; https://doi.org/10.5281/zenodo.17514848).

## Supporting information

### Supplementary Methods: Metabolomics

The gradient program for reverse-phase liquid chromatography (RPLC) was designed as follows: Chromatographic separation was performed using two distinct approaches with defined solvent systems. The mobile phases consisted of Solvent A (Water Lichrosolv® containing 0.1% formic acid) and Solvent B (Acetonitrile Honeywell® containing 0.1% formic acid). For RPLC, an Acquity UPLC BEH C18 column (1.7 µm, 2.1 × 150 mm) was used. The gradient started with 95% Solvent A for 1 minute, followed by a gradual decrease to 70% A over 14 minutes, then to 20% A within 2 minutes. This composition was held for 3 minutes, reduced further to 0.5% A in 2 minutes, maintained for 5 minutes, and finally returned to the initial 95% A within 4 minutes.

For hydrophilic interaction liquid chromatography (HILIC), separation was carried out on an Acquity UPLC BEH AMIDE column (1.7 µm, 2.1 × 100 mm). The program started at 5% Solvent A for 1 minute, increased to 30% A over 15 minutes, then to 80% A within 2 minutes, held for 1 minute, raised to 95% A within 1 minute, and finally returned to 4.5% A within 1 minute.

Both chromatographic methods were run under identical conditions: a flow rate of 0.4 mL/min, column temperature of 40 °C, and an injection volume of 5 µL. Mass spectrometry was performed after calibration with a mixture of 50 mL water, 50 mL 2-propanol, 1 mL NaOH, and 200 µL formic acid.

The mass spectrometer (MS) was operated in both positive and negative ionization modes, with the following settings: nebulizer pressure 2.0 bar, dry gas flow 8.0 L·min^− ?1^, dry gas temperature 200 °C, and capillary voltage of 4500 V (positive mode) or 3500 V (negative mode). The endplate offset was set at 500 V, and spectra were recorded across a mass range of 20–2000 m/z.

Metabolite identification was carried out using tandem mass spectrometry in auto MS/MS acquisition mode with smart exclusion. Precursor ions were selected when the difference between the rolling average and actual spectra exceeded 1000 counts. Fragmentation was achieved in the collision cell with collision energies ranging from 5 to 20 eV.

### Supplementary Tables

**Table S1:** Origin climatic information. Only environmental parameters relevant to drought tolerance in Q. robur and Q. petraea are shown (Nosenko et al., 2025).

|  | URB | NGL |
| --- | --- | --- |
| Latitude of the population origin site | 49,014 | 54,144 |
| Longitude of the population origin site | 8,156 | 13,072 |
| ID the German Weather Service (DWD) station nearest to the population origin site | 4177 | 6199 |
| Water holding capacity of soil in up to 1 m of effective rooting depth of vegetation period (mm) | 307,15 | 254,74 |
| Soil water available for plants in the effective rooting depth in the vegetation period (mm) | 170,13 | 137,24 |
| Mean annual precipitation (mm) | 803 | 657 |
| Mean annual temperature (°C) | 11,2 | 8,9 |
| Mean precipitation of vegetation period (mm) | 485 | 413 |
| Mean temperature of vegetation period (°C) | 16,9 | 14 |
| Mean precipitation of the driest month in Germany (mm) | 49 | 36 |

**Table S2:** Soil properties at the common garden (standard container substrate)

| Characteristic | per m<sup>3</sup> / value |
| --- | --- |
| white peat | 60 % |
| mixed peat | 20 % |
| wood fiber | 15 % |
| cocopeat | 5 % |
| wet clay | 75 kg |
| PG mix 14-16-18 (Yara, Brunsbüttel, Germany) | 1.2 kg |
| Radigen micronutrient slow-release fertilizer | 50 g |
| soil surfactant | 500 ml |
| pH target value | 5.8 |
| Structure | coarse |

**Table S3:** 16S read number during DADA2 pipeline.

| Sample | Input | Passed filter | DenoisedF | DenoisedR | Merged | Non chimeric | Passed total [%] |
| --- | --- | --- | --- | --- | --- | --- | --- |
| 1 | 95609 | 79096 | 72680 | 74535 | 61233 | 60932 | 63,73 |
| 2 | 103407 | 85008 | 79222 | 80490 | 69328 | 69045 | 66,77 |
| 3 | 114382 | 94742 | 87879 | 89367 | 74975 | 74574 | 65,2 |
| 4 | 105008 | 84055 | 78768 | 79850 | 68892 | 68386 | 65,12 |
| 5 | 118082 | 99291 | 91750 | 94084 | 78504 | 77750 | 65,84 |
| 6 | 99625 | 84657 | 77708 | 79635 | 65099 | 64443 | 64,69 |
| 7 | 90310 | 73307 | 68358 | 69332 | 59218 | 58824 | 65,14 |
| 8 | 80798 | 64409 | 59412 | 60305 | 49823 | 49626 | 61,42 |
| 9 | 137163 | 112692 | 105932 | 107552 | 93197 | 92773 | 67,64 |
| 10 | 132776 | 99073 | 93394 | 94415 | 79600 | 78860 | 59,39 |
| 11 | 106710 | 89889 | 82816 | 85269 | 70645 | 69757 | 65,37 |
| 12 | 69999 | 56571 | 52350 | 53121 | 44141 | 43883 | 62,69 |
| 13 | 84088 | 71021 | 65520 | 66838 | 55343 | 55155 | 65,59 |
| 14 | 80991 | 69038 | 62998 | 64728 | 52325 | 51915 | 64,1 |
| 15 | 61056 | 52030 | 46722 | 48702 | 38911 | 38827 | 63,59 |
| 16 | 83448 | 68637 | 63137 | 64906 | 53502 | 53069 | 63,6 |
| 17 | 79375 | 66479 | 60610 | 62750 | 50383 | 49874 | 62,83 |
| 18 | 59880 | 47719 | 43867 | 44798 | 37300 | 37172 | 62,08 |
| 19 | 67541 | 54602 | 50537 | 51258 | 42647 | 42591 | 63,06 |
| 20 | 143994 | 92847 | 86225 | 88485 | 73203 | 72751 | 50,52 |
| 21 | 70983 | 57683 | 53068 | 53897 | 44415 | 44272 | 62,37 |
| 22 | 134698 | 108752 | 101993 | 103045 | 88416 | 87461 | 64,93 |
| 23 | 123603 | 101229 | 95485 | 96806 | 84597 | 83715 | 67,73 |
| 24 | 93327 | 71744 | 66516 | 67505 | 55710 | 55608 | 59,58 |
| 25 | 98433 | 80319 | 74650 | 75518 | 64248 | 63903 | 64,92 |
| 26 | 104632 | 84761 | 78334 | 79469 | 67049 | 66633 | 63,68 |
| 27 | 143383 | 116422 | 110340 | 111245 | 98837 | 98139 | 68,45 |
| 28 | 76286 | 61227 | 56650 | 57400 | 47583 | 47455 | 62,21 |
| 29 | 409869 | 297612 | 284160 | 287540 | 253354 | 247768 | 60,45 |
| 30 | 103986 | 86442 | 81094 | 82338 | 72000 | 71459 | 68,72 |
| 31 | 118926 | 97542 | 91804 | 92602 | 80029 | 79649 | 66,97 |
| 32 | 132539 | 107807 | 100557 | 102156 | 86343 | 85969 | 64,86 |
| 33 | 127169 | 104181 | 98424 | 99283 | 87870 | 86842 | 68,29 |
| 34 | 104700 | 84829 | 78860 | 79886 | 67949 | 67621 | 64,59 |
| 35 | 105733 | 88195 | 82041 | 83386 | 70710 | 70363 | 66,55 |
| 36 | 73448 | 58457 | 54369 | 55499 | 47864 | 47663 | 64,89 |

**Table S4:**
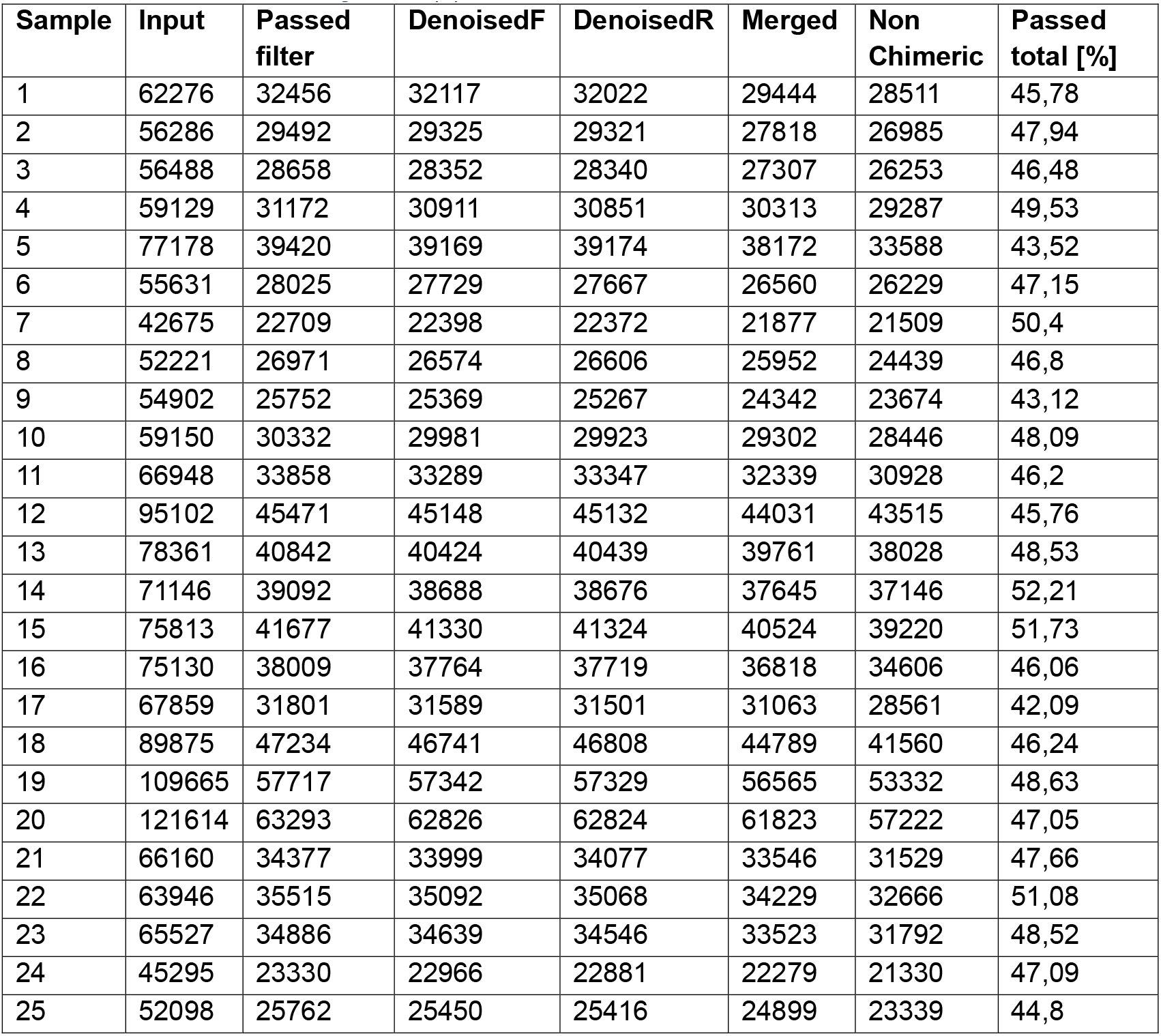
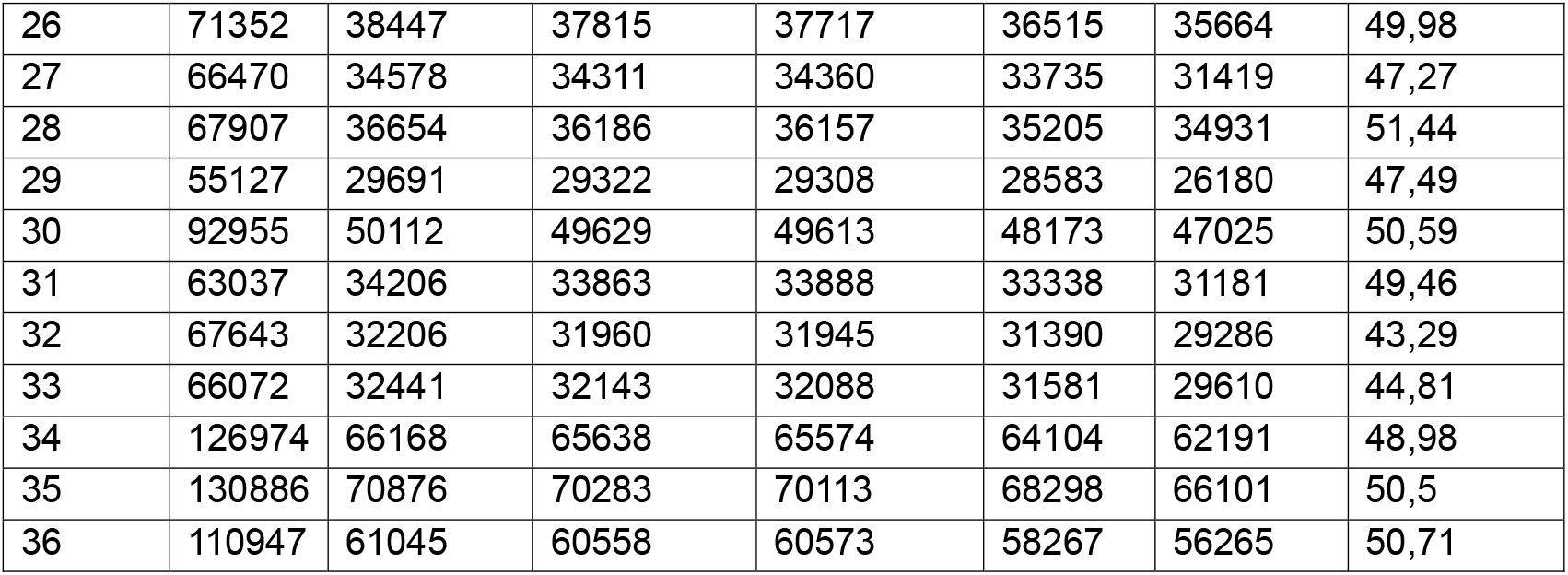
ITS reads lost during dada2 pipeline.

| Sample | Input | Passed filter | DenoisedF | DenoisedR | Merged | Non Chimeric | Passed total [%] |
| --- | --- | --- | --- | --- | --- | --- | --- |
| 1 | 62276 | 32456 | 32117 | 32022 | 29444 | 28511 | 45,78 |
| 2 | 56286 | 29492 | 29325 | 29321 | 27818 | 26985 | 47,94 |
| 3 | 56488 | 28658 | 28352 | 28340 | 27307 | 26253 | 46,48 |
| 4 | 59129 | 31172 | 30911 | 30851 | 30313 | 29287 | 49,53 |
| 5 | 77178 | 39420 | 39169 | 39174 | 38172 | 33588 | 43,52 |
| 6 | 55631 | 28025 | 27729 | 27667 | 26560 | 26229 | 47,15 |
| 7 | 42675 | 22709 | 22398 | 22372 | 21877 | 21509 | 50,4 |
| 8 | 52221 | 26971 | 26574 | 26606 | 25952 | 24439 | 46,8 |
| 9 | 54902 | 25752 | 25369 | 25267 | 24342 | 23674 | 43,12 |
| 10 | 59150 | 30332 | 29981 | 29923 | 29302 | 28446 | 48,09 |
| 11 | 66948 | 33858 | 33289 | 33347 | 32339 | 30928 | 46,2 |
| 12 | 95102 | 45471 | 45148 | 45132 | 44031 | 43515 | 45,76 |
| 13 | 78361 | 40842 | 40424 | 40439 | 39761 | 38028 | 48,53 |
| 14 | 71146 | 39092 | 38688 | 38676 | 37645 | 37146 | 52,21 |
| 15 | 75813 | 41677 | 41330 | 41324 | 40524 | 39220 | 51,73 |
| 16 | 75130 | 38009 | 37764 | 37719 | 36818 | 34606 | 46,06 |
| 17 | 67859 | 31801 | 31589 | 31501 | 31063 | 28561 | 42,09 |
| 18 | 89875 | 47234 | 46741 | 46808 | 44789 | 41560 | 46,24 |
| 19 | 109665 | 57717 | 57342 | 57329 | 56565 | 53332 | 48,63 |
| 20 | 121614 | 63293 | 62826 | 62824 | 61823 | 57222 | 47,05 |
| 21 | 66160 | 34377 | 33999 | 34077 | 33546 | 31529 | 47,66 |
| 22 | 63946 | 35515 | 35092 | 35068 | 34229 | 32666 | 51,08 |
| 23 | 65527 | 34886 | 34639 | 34546 | 33523 | 31792 | 48,52 |
| 24 | 45295 | 23330 | 22966 | 22881 | 22279 | 21330 | 47,09 |
| 25 | 52098 | 25762 | 25450 | 25416 | 24899 | 23339 | 44,8 |
| 26 | 71352 | 38447 | 37815 | 37717 | 36515 | 35664 | 49,98 |
| 27 | 66470 | 34578 | 34311 | 34360 | 33735 | 31419 | 47,27 |
| 28 | 67907 | 36654 | 36186 | 36157 | 35205 | 34931 | 51,44 |
| 29 | 55127 | 29691 | 29322 | 29308 | 28583 | 26180 | 47,49 |
| 30 | 92955 | 50112 | 49629 | 49613 | 48173 | 47025 | 50,59 |
| 31 | 63037 | 34206 | 33863 | 33888 | 33338 | 31181 | 49,46 |
| 32 | 67643 | 32206 | 31960 | 31945 | 31390 | 29286 | 43,29 |
| 33 | 66072 | 32441 | 32143 | 32088 | 31581 | 29610 | 44,81 |
| 34 | 126974 | 66168 | 65638 | 65574 | 64104 | 62191 | 48,98 |
| 35 | 130886 | 70876 | 70283 | 70113 | 68298 | 66101 | 50,5 |
| 36 | 110947 | 61045 | 60558 | 60573 | 58267 | 56265 | 50,71 |

**Table S5:** PERMANOVA results based on Bray-Curtis distances from microbiomes for factors origin, species, intrinsic water use efficiency (iWUE), and their interaction.

|  | **Prokaryotes** |  | **Fungi** |  |
| --- | --- | --- | --- | --- |
|  | **R<sup>2</sup>** | **p-value** | **R<sup>2</sup>** | **p-value** |
| Origin | 0.04 | 0.024* | 0.04 | 0.077 |
| Species | 0.03 | 0.662 | 0.03 | 0.493 |
| iWUE | 0.04 | 0.036* | 0.03 | 0.508 |
| Origin:Species | 0.03 | 0.468 | 0.04 | 0.105 |
| Origin:iWUE | 0.03 | 0.296 | 0.05 | 0.014* |
| Species:iWUE | 0.03 | 0.202 | 0.03 | 0.237 |
| Origin:Species:iWUE | 0.03 | 0.201 | 0.03 | 0.572 |

**Table S6:** Key metabolites identified through OPLS-DA with factor origin. Key metabolites have VIP score > 1, logFC > 1 and p-value <= 0.05)

| **Nr** | **VIP score** | **Log2FC** | **Adjusted p-value** | **Class** | **Assigned name** |
| --- | --- | --- | --- | --- | --- |
| 513 | 4.34 | -3.33 | 2.24E-04 | Protein | NA |
| 506 | 4.29 | -3.53 | 3.98E-05 | Not Matched | NA |
| 269 | 3.84 | 6.20 | 3.61E-03 | Protein | NA |
| 622 | 3.78 | -4.15 | 3.82E-04 | Lipid | NA |
| 444 | 3.60 | -3.84 | 5.69E-03 | Lipid | NA |
| 654 | 3.59 | -1.37 | 5.11E-03 | Lipid | NA |
| 508 | 3.55 | 3.81 | 5.11E-03 | Lipid | NA |
| 559 | 3.49 | -2.69 | 1.45E-03 | Oxyaromatic Compound | Ellagic Acid |
| 731 | 3.48 | -3.77 | 9.49E-04 | Protein | NA |
| 330 | 3.37 | -5.39 | 3.82E-04 | Lipid | NA |
| 87 | 3.32 | 5.35 | 3.52E-02 | Lipid | NA |
| 591 | 3.31 | -2.53 | 3.61E-03 | Not Matched | NA |
| 424 | 3.21 | 5.14 | 4.46E-03 | Protein | NA |
| 601 | 3.08 | -1.86 | 9.77E-03 | Protein | NA |
| 667 | 3.06 | -1.86 | 6.42E-03 | Carbohydrate | beta-Lactose |
| 437 | 3.04 | -2.40 | 1.45E-03 | Amino Sugar | NA |
| 170 | 3.00 | -5.71 | 1.26E-02 | Lipid | NA |
| 587 | 2.96 | -1.90 | 6.32E-03 | Carbohydrate | 1,6-Anhydro-β-D-glucose |
| 348 | 2.97 | -2.31 | 7.90E-04 | Protein | NA |
| 499 | 2.96 | -3.56 | 6.42E-03 | Protein | NA |
| 600 | 2.91 | -1.84 | 5.08E-03 | Protein | NA |
| 250 | 2.88 | -2.34 | 9.74E-03 | Not Matched | NA |
| 362 | 2.87 | -2.70 | 7.90E-04 | Protein | NA |
| 657 | 2.79 | -1.90 | 9.74E-03 | Carbohydrate | Raffinose |
| 413 | 2.77 | -2.39 | 5.11E-03 | Protein | NA |
| 304 | 2.75 | -2.86 | 1.45E-03 | Not Matched | NA |
| 399 | 2.68 | -3.08 | 3.82E-04 | Protein | NA |
| 684 | 2.64 | -1.45 | 2.51E-02 | Carbohydrate | NA |
| 398 | 2.56 | -2.07 | 5.82E-04 | Not Matched | NA |
| 255 | 2.53 | -2.65 | 2.29E-02 | Amino Sugar | NA |
| 168 | 2.32 | -2.50 | 4.34E-02 | Not Matched | NA |
| 419 | 2.25 | -2.14 | 1.75E-02 | Lipid | NA |
| 617 | 2.24 | -1.53 | 9.52E-03 | Protein | NA |
| 536 | 2.22 | -3.01 | 3.51E-02 | Oxyaromatic Compound | NA |
| 878 | 2.19 | -1.54 | 4.95E-02 | Amino Sugar | NA |
| 206 | 2.16 | -2.82 | 1.95E-02 | Oxyaromatic Compound | NA |
| 660 | 2.14 | -1.51 | 4.71E-02 | Protein | Arginine |
| 768 | 2.14 | -1.50 | 2.46E-02 | Oxyaromatic Compound | 3-Furoic acid |
| 507 | 2.12 | -1.10 | 8.32E-03 | Not Matched | NA |
| 504 | 2.10 | -1.33 | 2.72E-03 | Oxyaromatic Compound | NA |
| 311 | 2.10 | -1.72 | 2.84E-02 | Oxyaromatic Compound | NA |
| 361 | 2.08 | -1.23 | 1.09E-02 | Not Matched | NA |
| 400 | 2.04 | -1.62 | 4.86E-02 | Protein | NA |
| 471 | 2.03 | -1.92 | 4.33E-02 | Lipid | NA |
| 137 | 2.01 | -2.10 | 4.33E-02 | Oxyaromatic Compound | NA |
| 346 | 2.00 | -1.92 | 2.21E-02 | Lipid | NA |
| 750 | 1.20 | -1.28 | 2.51E-02 | Protein | NA |
| 913 | 1.98 | -1.55 | 3.06E-02 | Carbohydrate | NA |
| 324 | 1.95 | -2.18 | 2.69E-02 | Lipid | NA |
| 358 | 1.89 | 1.58 | 1.67E-02 | Not Matched | NA |
| 883 | 1.76 | -1.07 | 1.40E-02 | Oxyaromatic Compound | NA |
| 179 | 1.75 | -1.02 | 4.33E-02 | Amino Sugar | NA |
| 25 | 1.73 | -1.41 | 3.65E-02 | Lipid | NA |
| 120 | 1.72 | -1.12 | 2.15E-02 | Carbohydrate | NA |
| 301 | 1.70 | -1.29 | 4.34E-02 | Oxyaromatic Compound | NA |
| 1000 | 1.70 | -1.10 | 4.00E-02 | Oxyaromatic Compound | NA |
| 153 | 1.63 | -1.47 | 4.58E-03 | Not Matched | NA |
| 734 | 1.61 | -1.11 | 2.51E-02 | Not Matched | NA |

### Supplementary Figures

**Figure S1:**
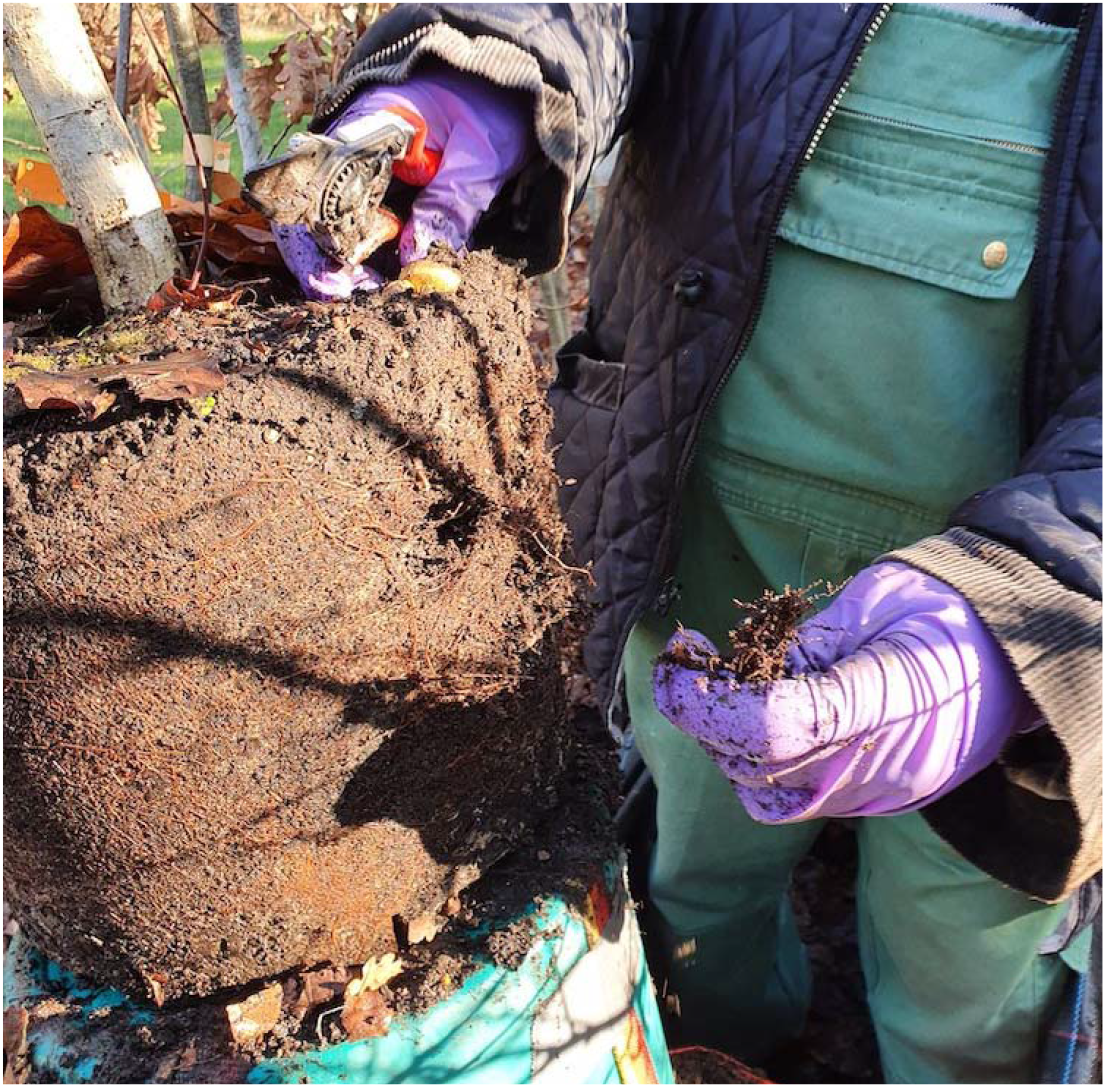
Illustration of the rhizosphere sampling procedure (February 2023)

**Figure S2:**
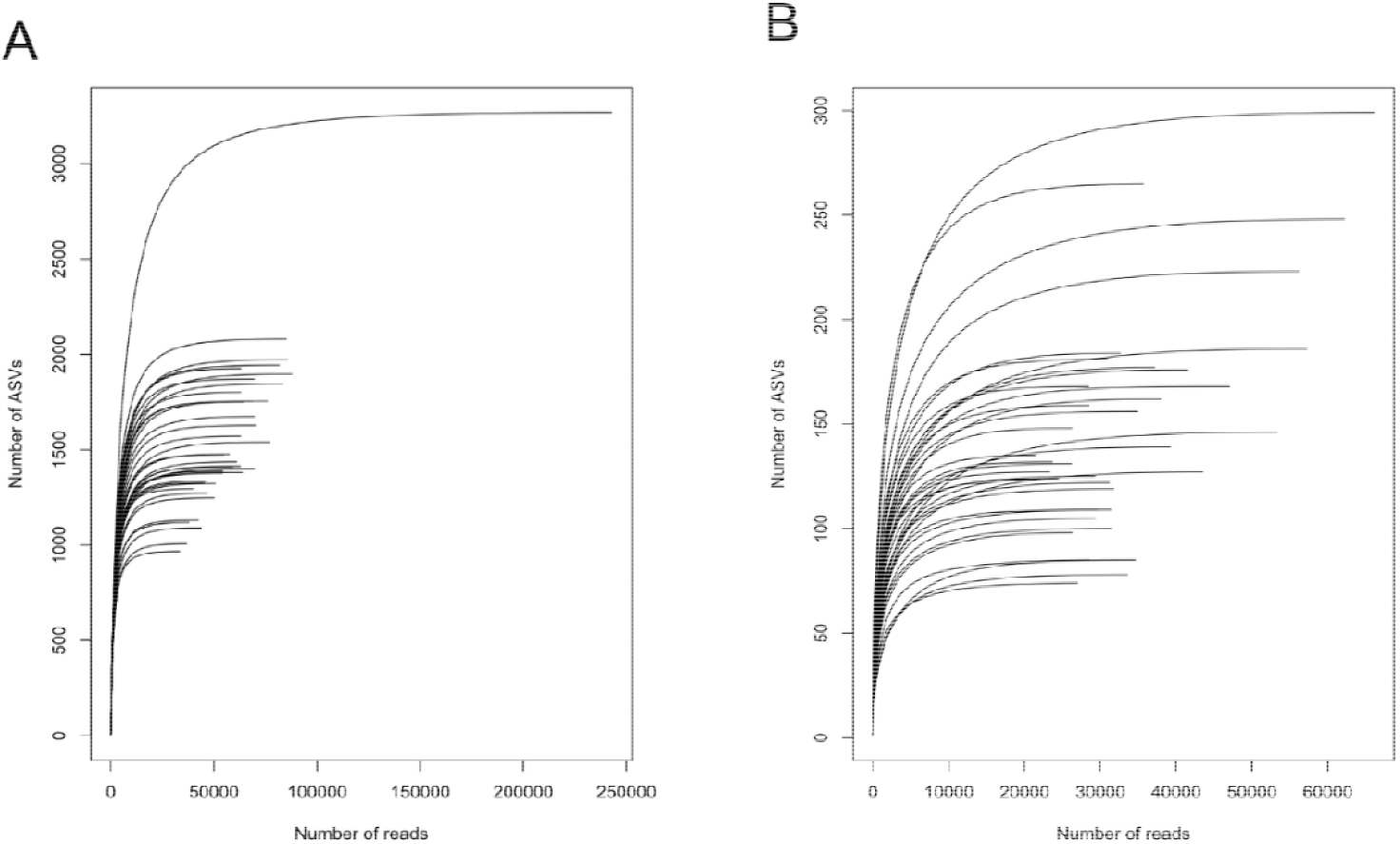
Rarefaction curves for 16S (A) and ITS (B) sequencings drawn from unnormalized abundance table resulting from the dada2 pipeline

**Figure S3:**
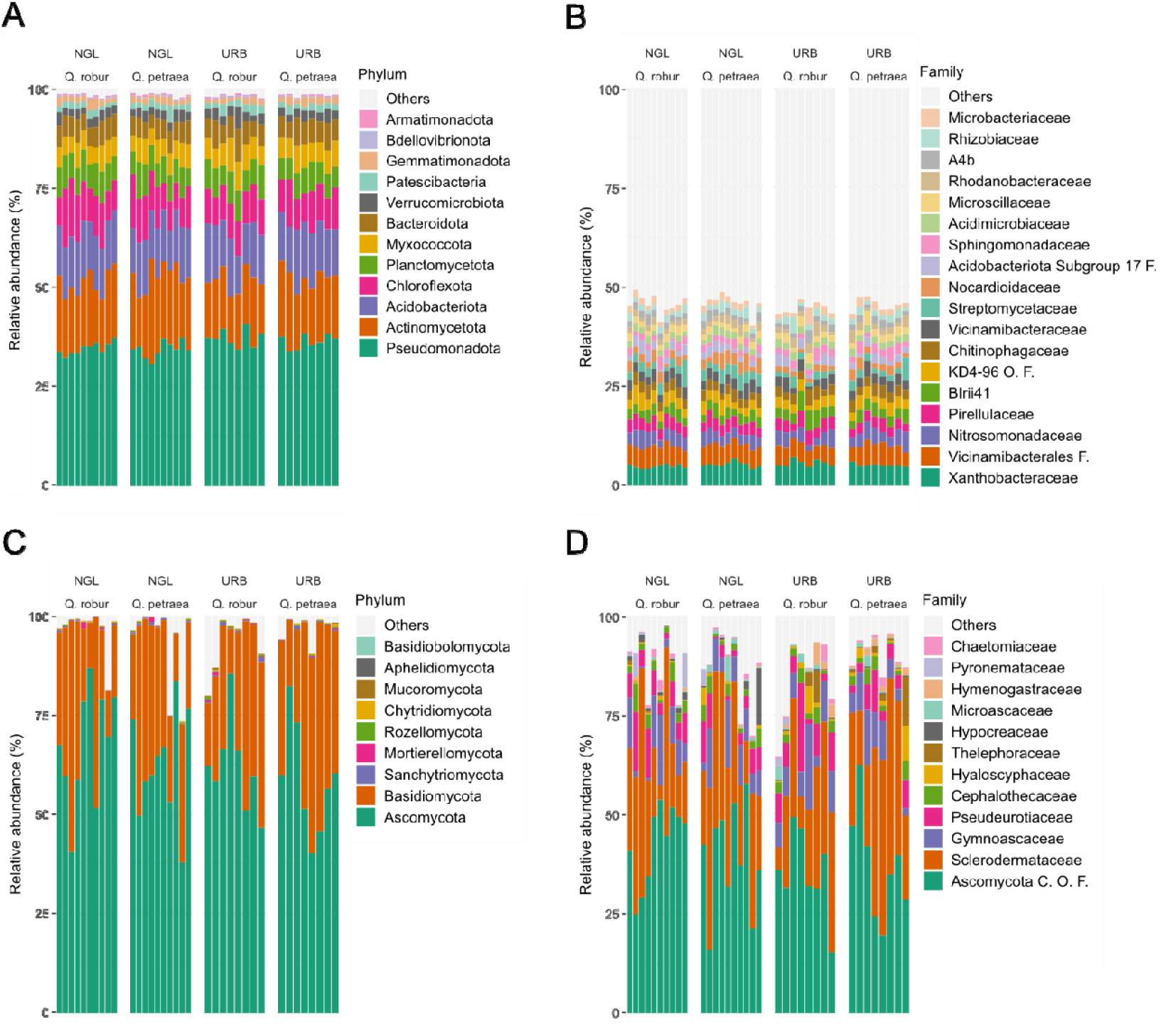
Bacterial and fungal community composition at phylum and family level. Shown are the relative abundance of the most prominent prokaryotic phyla (A), bacterial families (B), fungal phyla (C), and fungal families (D).

**Figure S4:**
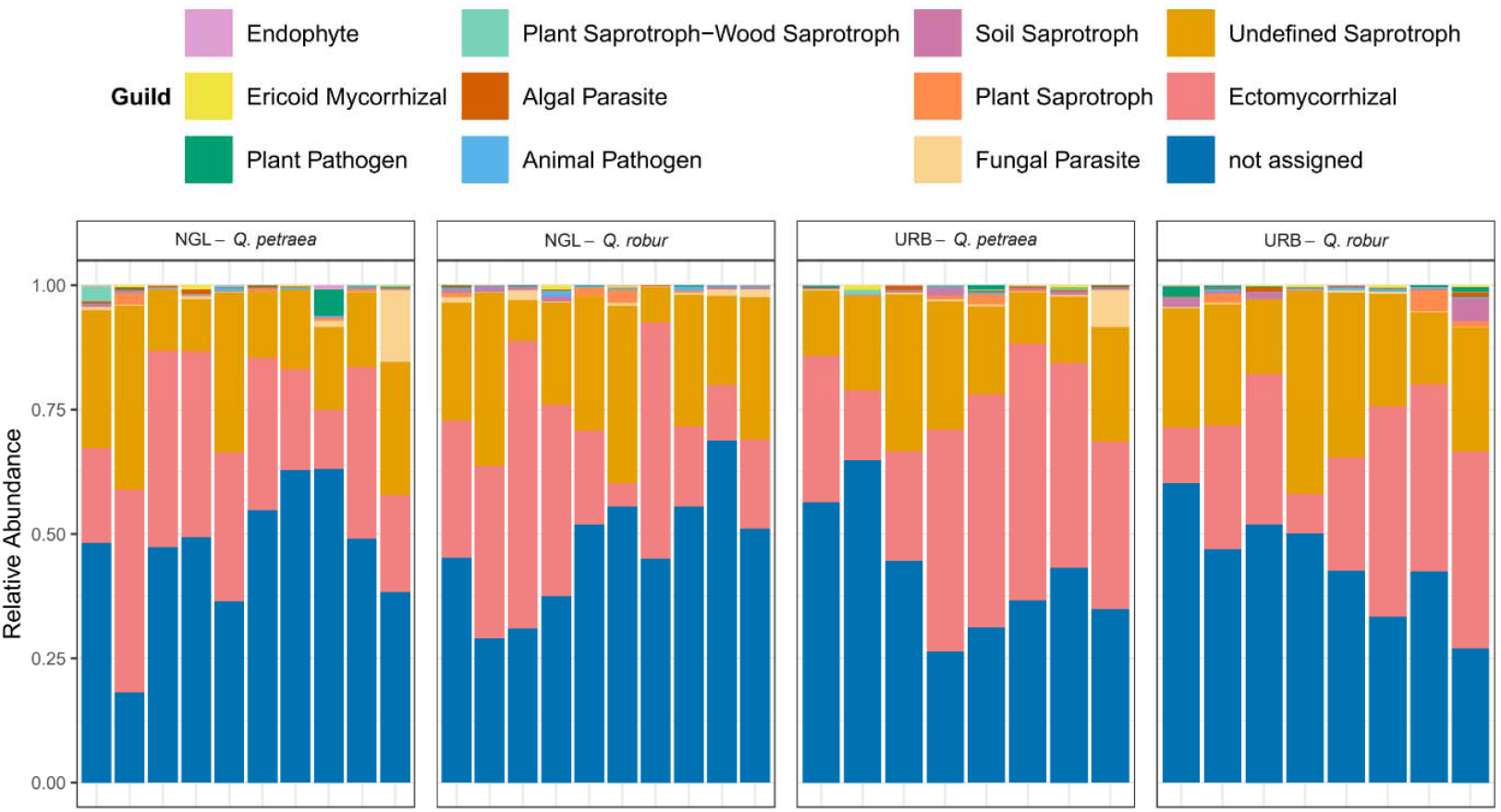
Fungal guild classification of ITS reads per sample. Classification was performed using FunGuild. Guilds with a total read amount of less than 1000 were excluded from visualization. The guild assignment with the highest likelihood was chosen for each ASV.

**Figure S5:**
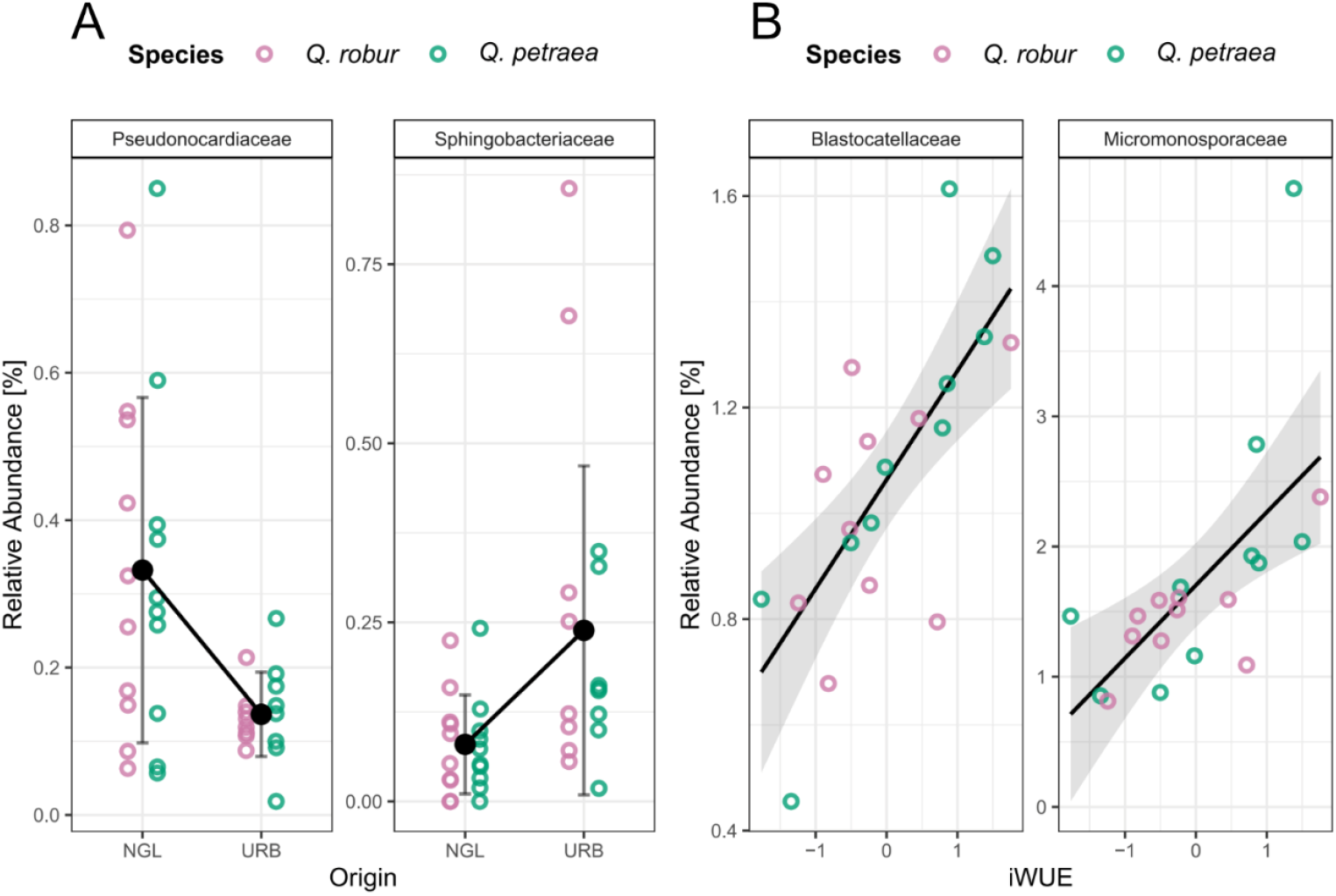
Differentially abundant prokaryotic families found thought DESeq2 analysis. (A) Families associated with one of the origins. (B) Families associated with intrinsic water use efficiency (iWUE).

**Figure S6:**
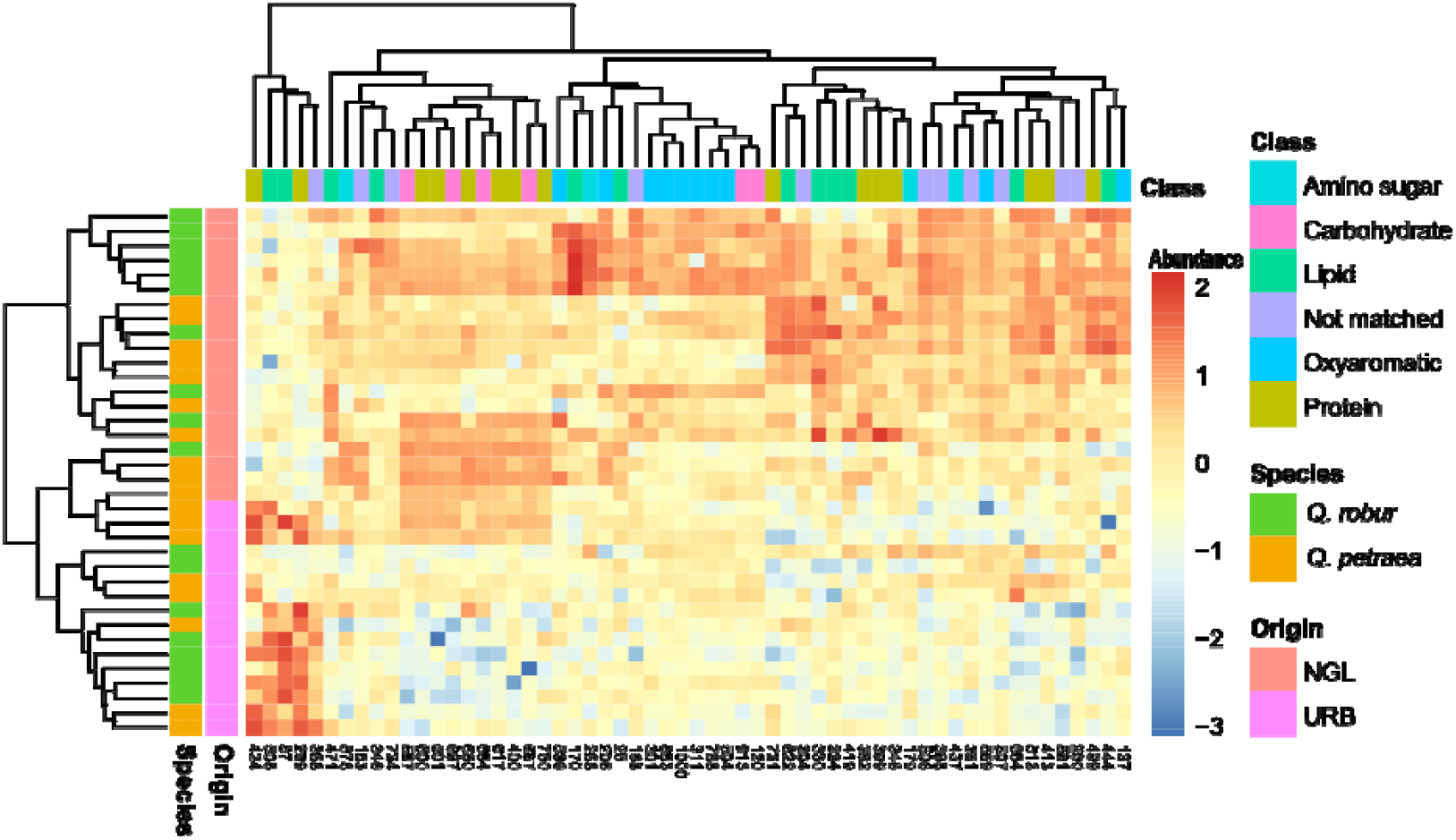
Abundance of key metabolites identified through OPLS-DA (VIP>1, log-FC > 1, p-value <= 0.05). Metabolite data was log10-transformed and Pareto-scaled.

**Figure S7:**
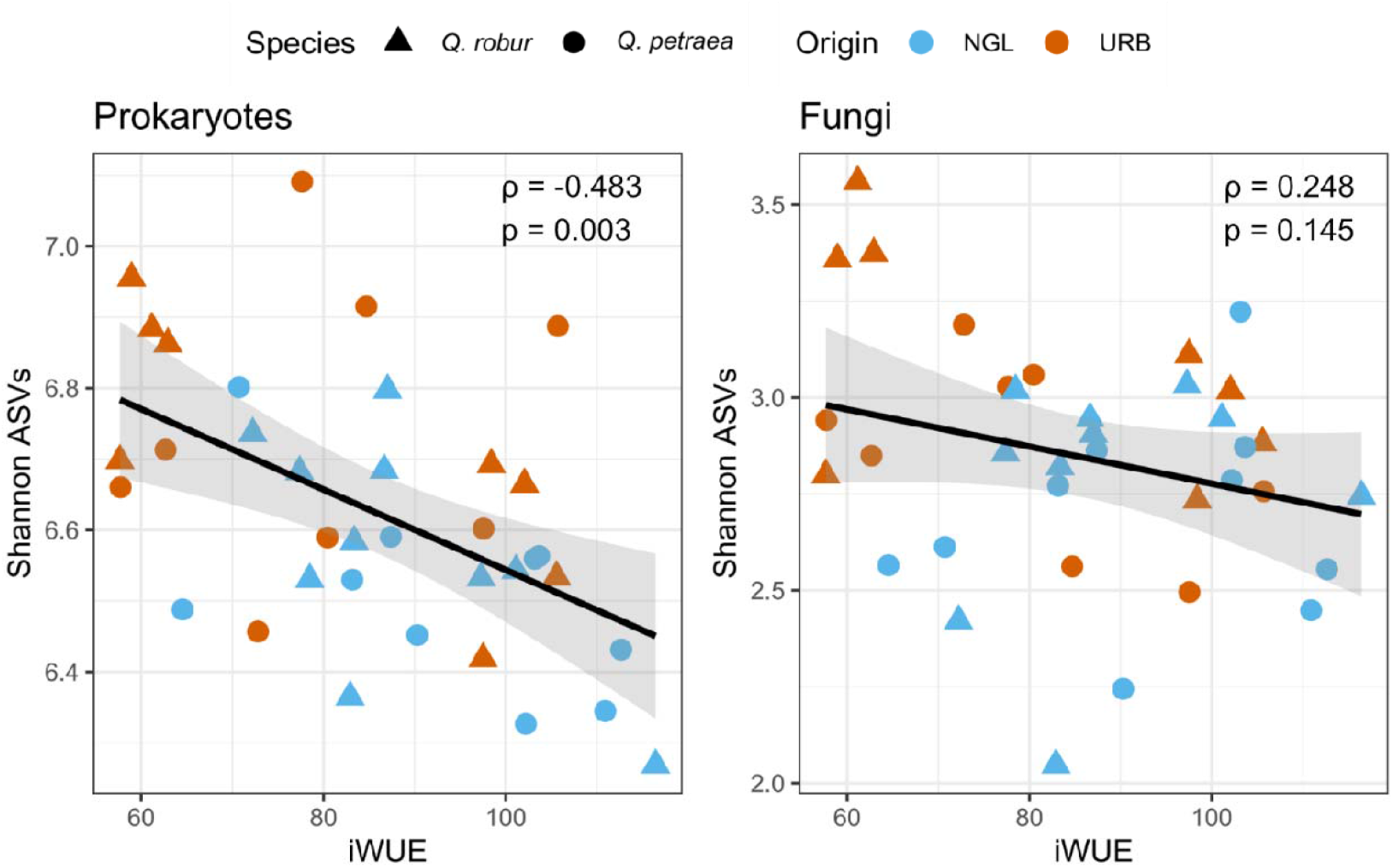
Spearman correlation of microbial Shannon diversity and intrinsic water use efficiency (iWUE) across origins and sites. Spearman ρ and p-value are indicated.

**Figure S8:**
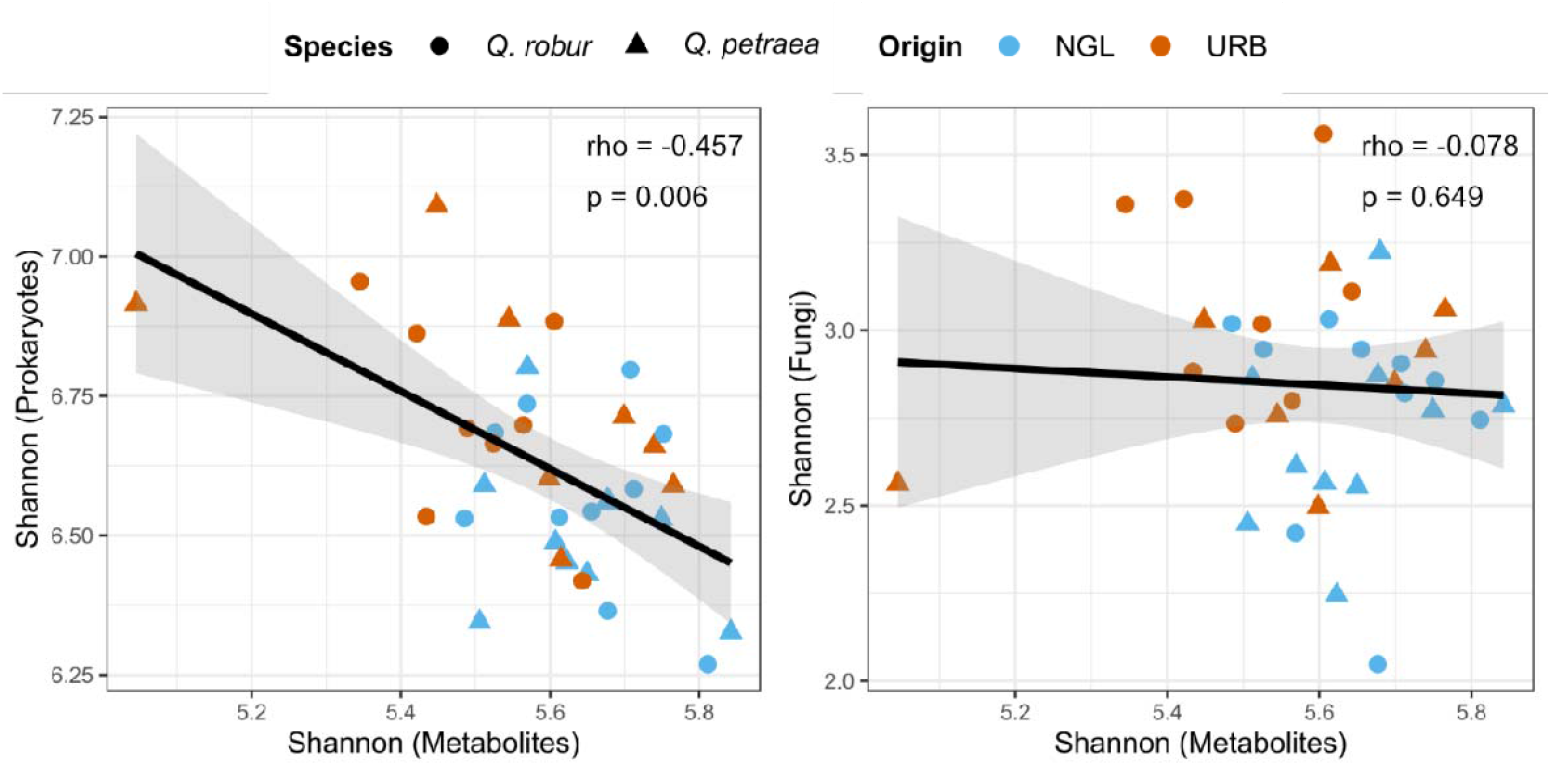
Correlation of metabolite Shannon diversity with prokaryotic (left) and fungal (right) Shannon diversity indices. Correlation of prokaryotic microbiome and metabolome diversity was also significant for the URB data subset (rho = −0.57, p = 0.0249). Values denoted in the plot refer to whole dataset tests

## Notes

### Competing Interest Statement

The authors have declared no competing interest.

